# Quantifying the effects of single mutations on viral escape from broad and narrow antibodies to an H1 influenza hemagglutinin

**DOI:** 10.1101/210468

**Authors:** Michael B. Doud, Juhye M. Lee, Jesse D. Bloom

**Affiliations:** Basic Sciences and Computational Biology Program, Fred Hutchinson Cancer Research Center; Department of Genome Sciences, University of Washington Seattle, WA, USA; Medical Scientist Training Program, University of Washington Seattle, WA, USA

## Abstract

Influenza virus can completely escape most antibodies with single mutations. However, rare antibodies broadly neutralize many viral strains. It is unclear how easily influenza virus might escape such antibodies if it was under strong pressure to do so. Here we map all single amino-acid mutations that increase resistance to broad antibodies targeting an H1 hemagglutinin. Crucially, our approach not only identifies antigenic mutations but also quantifies their effect sizes. All antibodies select mutations, but the effect sizes vary widely. The virus can escape a broad antibody that targets residues in hemagglutinin’s receptor-binding site the same way it escapes narrow strain-specific antibodies: via single mutations with huge effects. In contrast, broad antibodies targeting hemagglutinin’s stalk only select mutations with small effects. Therefore, among the antibodies we have examined, breadth is an imperfect indicator of the potential for viral escape via single mutations. Broadly neutralizing antibodies targeting the H1 hemagglutinin stalk are quantifiably harder to escape than the other antibodies tested here.

## INTRODUCTION

Nearly all viruses show some antigenic variation. However, the extent of this variation ranges widely. For instance, although both measles virus^1,2^ and polio virus^3,4,5^ exhibit antigenic variation, the magnitude of this variation is small. Therefore, immunity to these viruses is lifelong^6,7^. In contrast, human influenza virus exhibits much more antigenic variation. So although infection with an influenza virus strain provides long-term immunity to that exact strain^8,9,10^, the virus’s rapid antigenic evolution erodes the effectiveness of this immunity to that strain’s descendants within ~5 years^11,12^.

One possible reason that viruses exhibit different amounts of antigenic variation is that they have disparate evolutionary capacities to escape the immunodominant antibodies generated by natural immune responses^13,14,15^. According to this explanation, human influenza virus undergoes rapid antigenic drift because most neutralizing antibodies target epitopes on the viral hemagglutinin (HA) protein that are highly tolerant of mutational change. This explanation is supported by classic experiments showing that it is easy to select viral mutants that escape most antibodies^16,17^, as well as by the observation that mutations that alter antigenicity arise frequently during influenza’s evolution globally^18,19,20,21,22^ and within individual humans with long-term infections^23^. A corollary of this explanation is that influenza virus’s capacity for antigenic drift would be reduced if most antibodies instead targeted epitopes that were less mutationally tolerant.

Verifying this corollary has become of practical importance with the discovery of broadly neutralizing antibodies against influenza virus. These antibodies typically target conserved epitopes in HA’s stalk^24,25,26^ or receptor-binding site^27,28,29^, and neutralize a wide range of viral strains. Broad antibodies are usually less abundant in human serum than antibodies to antigenically variable epitopes on the head of HA^30,31^. However, major efforts are underway to elicit broad antibodies by vaccination or administer them directly as therapeutics^32,33^.

If these efforts succeed, the epitopes of broad antibodies could come under stronger antigenic selection in human influenza virus. Might such selection then drive antigenic variation in these epitopes? There is precedent for the idea that the immune status of the host population can shape influenza virus evolution: the virus undergoes faster antigenic drift in long-lived humans that accumulate immune memory than in short-lived swine that are mostly naive^34,35^, and poultry vaccination may accelerate antigenic drift of avian influenza^36,37^. But alternatively, perhaps broad antibodies are broad because the virus has difficulty escaping them regardless of selection from host immunity.

So far, there is limited data to distinguish between these possibilities. Several studies have shown that the head domain of HA is more mutationally tolerant than the stalk domain where many broad antibodies bind^38,39,40^. However, these studies did not select for antibody escape, so it is difficult to relate their measurements to the virus’s evolutionary capacity under immune selection. Other work has shown that it is possible to select antigenic mutants with broad antibodies^41,42,43,44,45,46^, demonstrating that these epitopes are not entirely refractory to change. But given that antibodies can select some antigenic variation even in measles virus^1,2^ and polio virus^3,4^, the existence of selectable mutations does not necessarily imply that influenza virus can escape broad antibodies as easily as it drifts away from narrow strain-specific ones. The fundamental problem is that existing studies have not quantified the ease of viral escape in a way that can be compared across antibodies in an apples-to-apples fashion.

Here we systematically quantify the results of selecting all single amino-acid mutations to an H1 HA with several broad and narrow antibodies. Critically, our approach quantifies the *magnitude* of the antigenic effect of every mutation in a way that can be directly compared across antibodies. We find that even the broadest antibodies select antigenic mutations. However, the magnitudes of the antigenic effects vary greatly across antibodies. Single mutations make the virus completely resistant to both narrow strain-specific antibodies and a broad antibody that targets residues in HA’s receptor-binding site. But no single mutation does more than modestly increase the virus’s resistance to two broad antibodies against the HA stalk. Therefore, broad anti-stalk antibodies are quantifiably more resistant to viral escape via single amino-acid mutations than the other antibodies tested here.

## RESULTS

### An approach to quantify the fraction of virions with each mutation that escape antibody neutralization

We can visualize the outcome of antibody selection on viral populations containing antigenic mutations as in Figure 1. If a mutation strongly escapes neutralization, then all virions with this mutation survive antibody treatment at a concentration where other virions are mostly neutralized (Figure 1A). This escape is manifested by a large shift in the neutralization curve for the mutant (Figure 1B). If we draw vertical lines through the overlaid neutralization curves, we can calculate the fraction of virions with each mutation that survive neutralization at each antibody concentration. These fractions can be represented using logo plots, where the height of each letter is proportional to the fraction of virions with that amino acid at a site that survive (Figure 1C). Large letters correspond to strong escape mutations.

**Figure 1:**
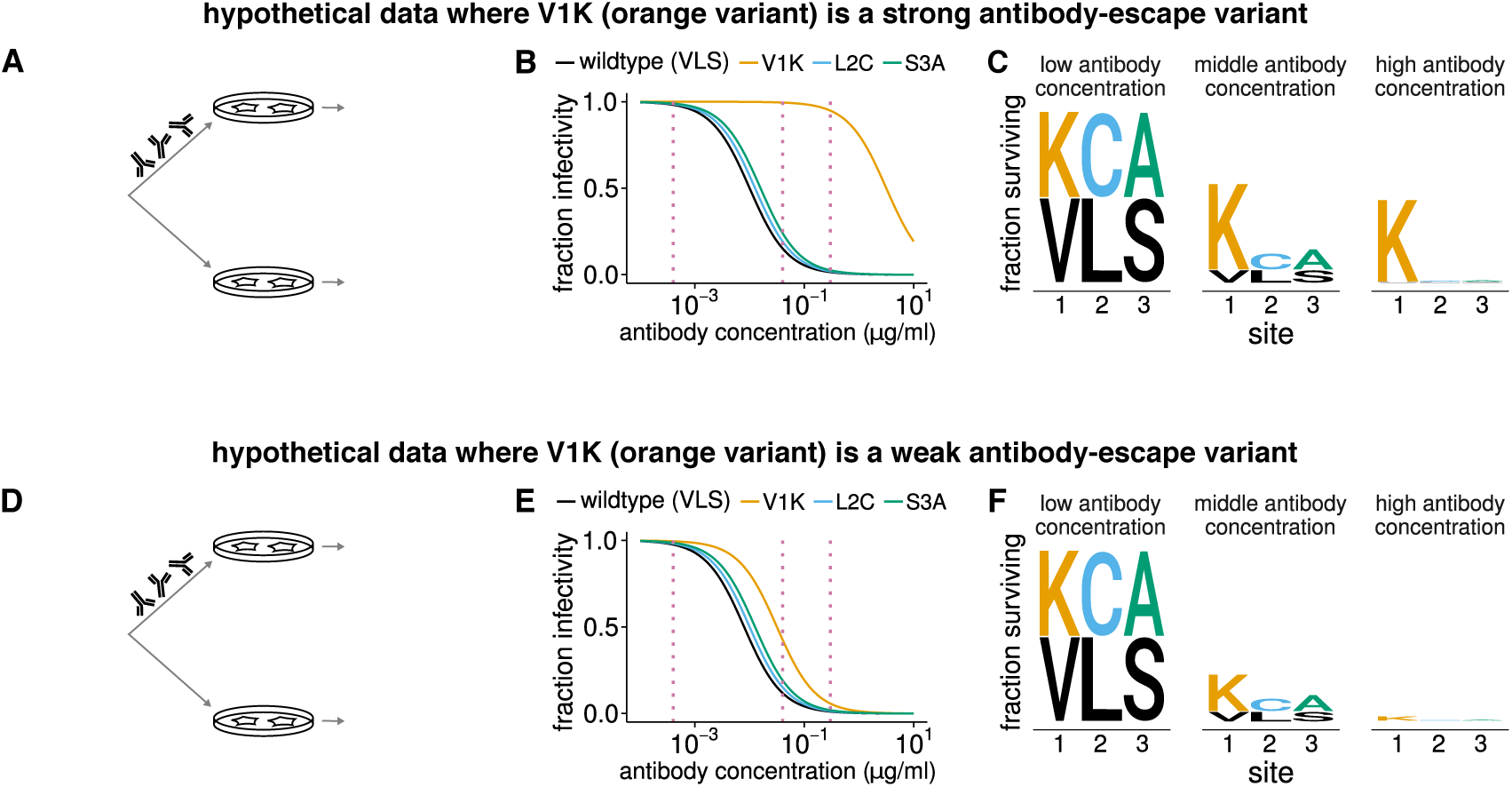
Quantifying the fraction of virions with each mutation that escape antibody neutralization. This figure shows hypothetical data for four viral variants. (A) Virions with the V1K mutation (orange) completely survive an antibody concentration where most other virions are neutralized. (B) This resistance is manifested by a large shift in V1K’s neutralization curve. (C) For each dotted vertical line drawn through the neutralization curves in (B), we calculate the fraction of virions with that mutation that survive the antibody, and indicate this fraction by the height of the letter corresponding to that amino acid at that site. (D-F) Similar data to the first three panels, but now V1K has only a small antigenic effect, and so only modestly increases the fraction of virions that survive antibody treatment.

Now consider the case where a mutation has just a small antigenic effect, and so only slightly increases the fraction of virions that survive neutralization (Figure 1D). In this scenario, the neutralization curve shifts only slightly (Figure 1E). In the logo plot representation, the antigenic mutation is only slightly larger than other amino acids (Figure 1F), since possessing the mutation only modestly increases the chance that a virion survives antibody treatment. These logo plots therefore provide a way to both identify antigenic mutations and quantify the magnitudes of their effects in a way that is directly comparable across antibodies.

Our goal is to determine the fraction of mutant virions that survive antibody neutralization for *all* mutations to HA. One way to do this would be to measure individual neutralization curves for each of the 19 × 565 = 10,735 single amino-acid mutants of the 565-residue HA protein. However, individually creating and assaying that many mutants would be exceedingly time-consuming and expensive. Fortunately, we have shown that antibody selection on all viral mutations can be assayed in a single experiment using mutational antigenic profiling^47,48^. This approach involves generat-ing viral libraries containing all mutations to the protein of interest, selecting these viruses with or without antibody, and using an accurate deep-sequencing method to determine the relative frequencies of each mutation.

These frequencies can be analyzed to calculate the fraction of virions with each mutation that survive antibody treatment. Specifically the deep sequencing determines the frequencies of virions carrying amino-acid *a* at site *r* in the antibody-selected and mock-selected conditions, which we denote as 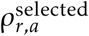 and 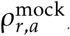, respectively. We can also measure the total fraction of the viral library that survives the antibody which we denote as *γ* The fraction of variants with amino-acid *a* at site *r* that survive antibody selection is then simply

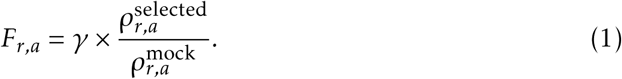

For instance, in Figure 1A, the frequency of virions with the orange mutation is 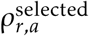 = 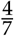 in the antibody selection and 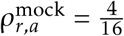 in the mock selection. The overall fraction of virions that survive the antibody in Figure 1A is 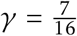. Therefore, we use Equation 1 to calculate that the fraction of variants with the orange mutation that survive is 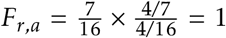. Performing the analogous calculation for Figure 1D correctly determines that fraction of virions with the orange mutation that survive the antibody is only 0.5 for the scenario in that figure panel. In the analyses of real data below, we will plot the excess fraction surviving *above* the overall library average, which is

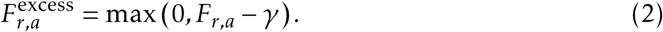

Importantly, Equations 1 and 2 correct for effects on viral growth due to normalization by the mock-selected control, and so measure only antigenicity and not viral growth provided that the virus at least grows well enough to be present in the library. Details of how the calculations are extended to account for sequencing errors and sampling statistics are in the METHODS. Open-source software that performs all steps in the analysis beginning with the deep sequencing data is available at https://jbloomlab.github.io/dms_tools2/.

### Broad and narrow antibodies that neutralize influenza virus

We applied this approach to anti-HA antibodies with a range of breadths and epitopes. The crystal structures or sites of escape mutations selected by these antibodies are shown in Figure 2A. We chose two broad antibodies, FI6v3 and C179, that target the stalk of HA^26,51,50^. FI6v3 is extremely broad, and neutralizes both group 1 and group 2 HAs (Figure 2B). C179 is less broad, and neutralizes only some group 1 HAs (Figure 2B). We also chose a broad antibody, S139/1, that crystallographic studies have shown binds to residues in HA’s receptor-binding pocket^27^, and which can neutralize both group 1 and group 2 HAs^41,27^. Finally, we re-analyzed deep sequencing data from prior mutational antigenic profiling of three narrow strain-specific antibodies, H17-L19, H17-L10, and H17-L7^47^. These narrow antibodies bind the Ca2, Ca1, and Cb antigenic regions on HA’s globular head^52^, and only neutralize a narrow slice of H1 viruses.

**Figure 2:**
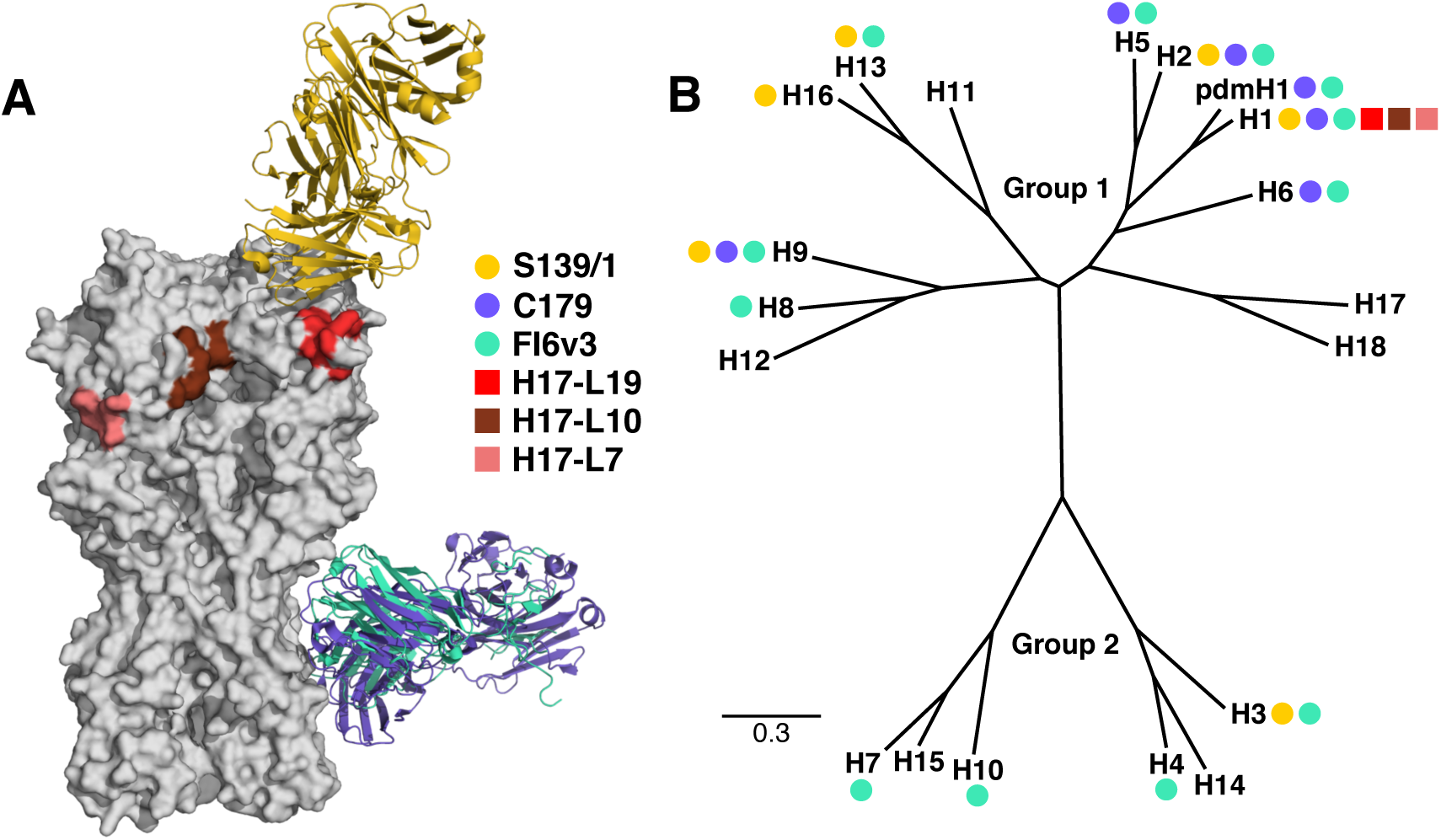
Epitopes and breadth of broad and narrow antibodies targeting HA. (A) Crystal structures of the broad antibodies and sites of escape mutations selected by the narrow ones superimposed on the structure of the HA trimer (PDB 1RVX^49^). S139/1 (PDB 4GMS^27^) targets residues in the receptor-binding pocket; C179 (PDB 4HLZ^50^) and FI6v3 (PDB 3ZTN^26^) target the stalk. The sites of escape mutations for H17-L19, H17-L10, and H17-L7 are those mapped by Doud et al^47^. (B) A phylogenetic tree of HA subtypes. Circles (broad antibodies) and squares (narrow antibodies) denote reported antibody binding or neutralization activity against that subtype. Not all antibodies have been tested against all subtypes.

We performed our experiments using the lab-adapted A/WSN/1933 (H1N1) strain of influenza. This strain is derived from an early seasonal H1N1 that was extensively passaged in the lab, where it adapted to become neurotropic and trypsin in-dependent^53^. But despite these unusual properties, the virus is neutralized by most broad antibodies that target other H1 viruses, including those used in this study (Figure 3). Our experiments utilize fully infectious influenza virus rather than pseudovirus, which is important since the accessibility of some epitopes can vary with HA density, which differs between fully infectious virus and pseudovirus^26,54^.

**Figure 3:**
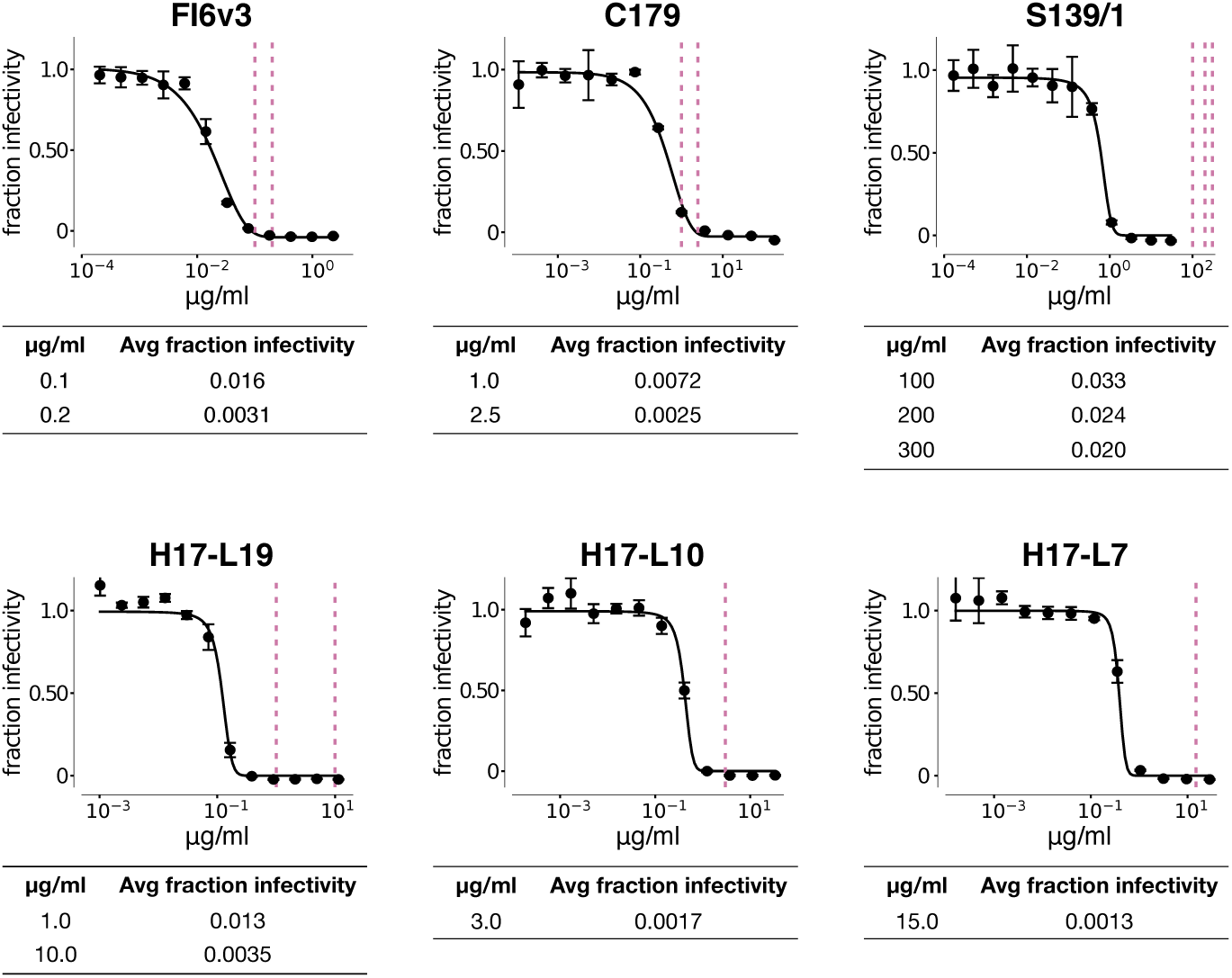
Neutralization of wildtype virus by each antibody, and the fraction of mutant library virions surviving at each concentration used in our experiments. The curves show neutralization of the wildtype A/WSN/1933 virus. Each point represents the mean and standard deviation of three measurements. The vertical dotted lines show the concentrations of antibody that were then used in the mutant virus library selections, and the tables give the overall fraction of the mutant virus libraries that survived at each concentration, determined by qRT-PCR. As described in the text, the antibody concentrations were chosen to give similar fractions of the mutant virus libraries that survive, rather than to fall at uniform positions on the neutralization curves of the wildtype virus.

The wildtype virus is neutralized by all the antibodies, with IC50s between 0.01 and 1 *μ*g/ml (Figure 3). However, our selections are performed on mutant virus libraries, not wildtype virus. Because these libraries have different capacities to escape each antibody, the fraction of each library that survives high antibody concentrations will vary among antibodies. For instance, at concentrations that neutralize 99% of the wildtype virus, we expect a larger fraction of a library to survive an antibody for which there are many HA escape mutations than an antibody with few HA escape mutations. Therefore, rather than using the same concentration for all antibodies, we selected concentrations for each antibody where between 2% and 0.1% of the libraries survived in order to strongly select for escape mutations (Figure 3). Slight differences among antibodies in the fraction surviving within this range should not strongly affect our results, since Equations 1 and 2 account for such differences via the *γ* term. However, to confirm the robustness of our results, we used several concentrations of each broad antibody (Figure 3).

### Quantifying the antigenic effects of all mutations selected by each antibody

We performed mutational antigenic profiling using the three broad antibodies at the concentrations indicated in Figure 3. All experiments were performed in full biological triplicate using three independently generated virus libraries carrying single amino-acid mutations to HA^55^. Importantly, as described previously^55^, these virus libraries were generated by mutagenizing HA at the *codon* level rather than at the nucleotide level. Performing codon mutagenesis is important, because single-nucleotide mutations access only about a third of the possible amino-acid mutations from a given codon, whereas codon mutations access all possible amino-acid mutations.

The correlations among replicates of the mutational antigenic profiling, in terms of the measured fraction-surviving above average for each possible amino-acid mutation, are shown in Figure S1. For the remainder of this paper, we will refer to the median antigenic effect of each mutation across replicates.

It is immediately obvious that the narrow strain-specific antibodies and the antibody targeting residues in HA’s receptor-binding pocket (S139/1) select mutations with large antigenic effects. For all four of these antibodies, there are multiple sites in HA where mutations enable a substantial fraction of virions to survive high antibody concentrations (Figure 4). Specifically, there are mutations that enable over a third of virions to survive at concentrations where virtually all wildtype virions are neutralized (Figure S2). Therefore, the virus can escape these four antibodies with the sort of large-effect single amino-acid mutations that characterize traditional influenza antigenic drift^16,^17^,18,19,20,21^.

**Figure 4:**
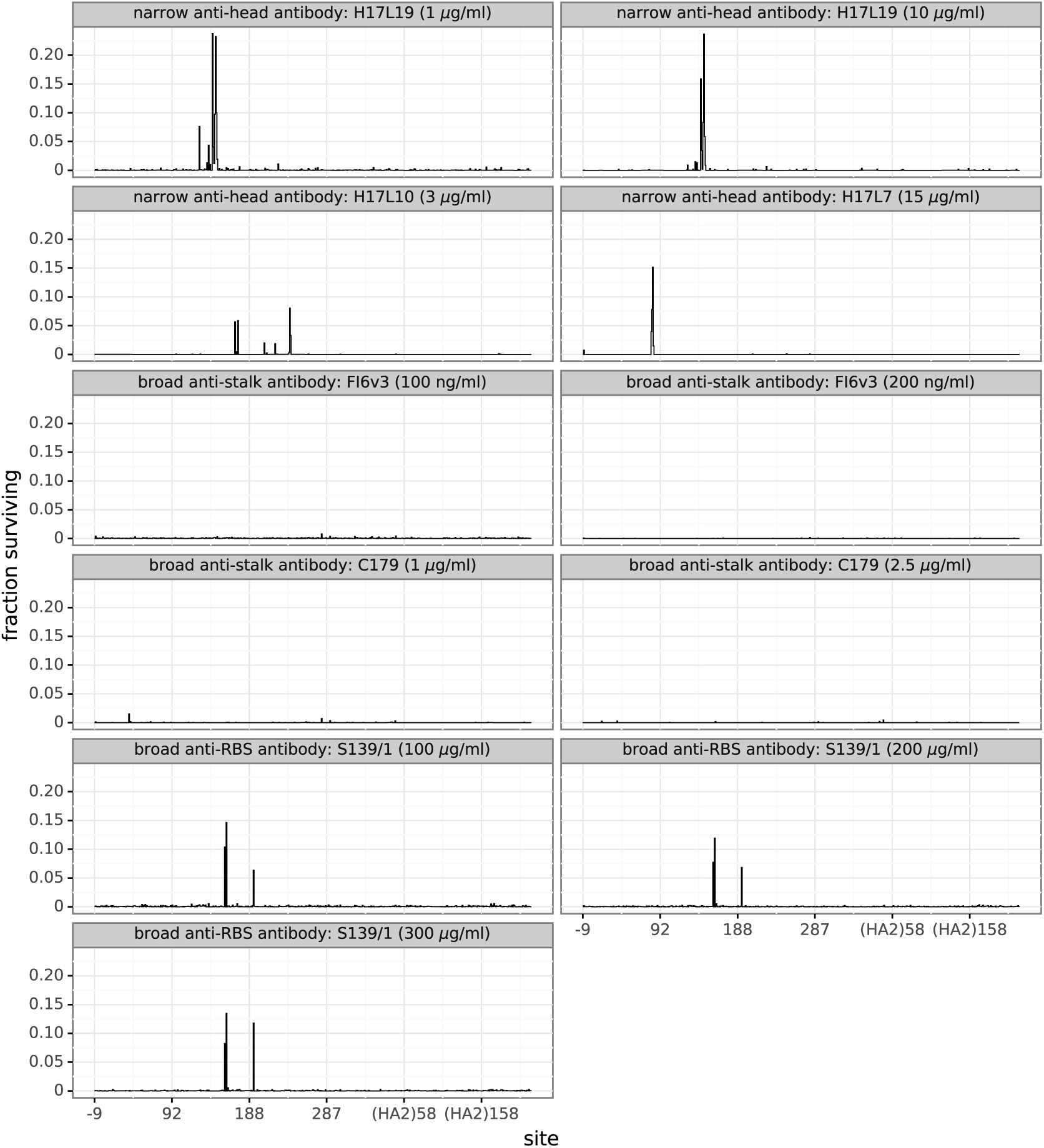
Strain-specific and anti-receptor-binding-site antibodies select mutations with large antigenic effects, but anti-stalk antibodies only select small-effect mutations. The excess fraction of virions with a mutation at each site that survive the antibody, averaging across all amino-acid mutations at each site (see Equation 10). There are multiple sites of large-effect mutations for H17L19, H17L10, H17L7, and S139/1—but none for FI6v3 and C179. Figure S2 shows the excess fraction surviving for the largest-effect mutation at each site. Figures S3, S4, S5, S6, S7, and S8 show all mutations using logo plots. Sites are labeled in H3 numbering.

In contrast, the stalk-targeting antibodies C179 and FI6v3 select no strong escape mutants. If we look at the results for these antibodies on the same scale as the other antibodies, we see only a few small bumps in the fraction of virions surviving (Figures 4 and S2). Only if we zoom in can we see that there are actually a few sites where mutations slightly increase the fraction of virions surviving C179 and FI6v3 (Figures S3, S4, S5, S6, S7, and S8). But the effect sizes of these antigenic mutations are tiny compared to the other antibodies—especially for FI6v3. Therefore, the HA of A/WSN/1933 influenza virus is far less capable of escaping these anti-stalk antibodies by single mutations than it is of escaping the other four antibodies.

### The selected mutations are near the binding footprints of the antibodies

Antigenic mutations selected by narrow strain-specific antibodies against HA are thought to occur at residues in or near the physical binding footprint of the antibody^16,17,52^. We examined whether this was the case for the broad antibodies used in our experiments. Figure 5A shows a zoomed-in view of the sites of mutations selected by each antibody, as well as their locations on HA’s structure. It is immediately clear that the selected mutations are nearly all in or close to the antibody-binding footprint.

**Figure 5:**
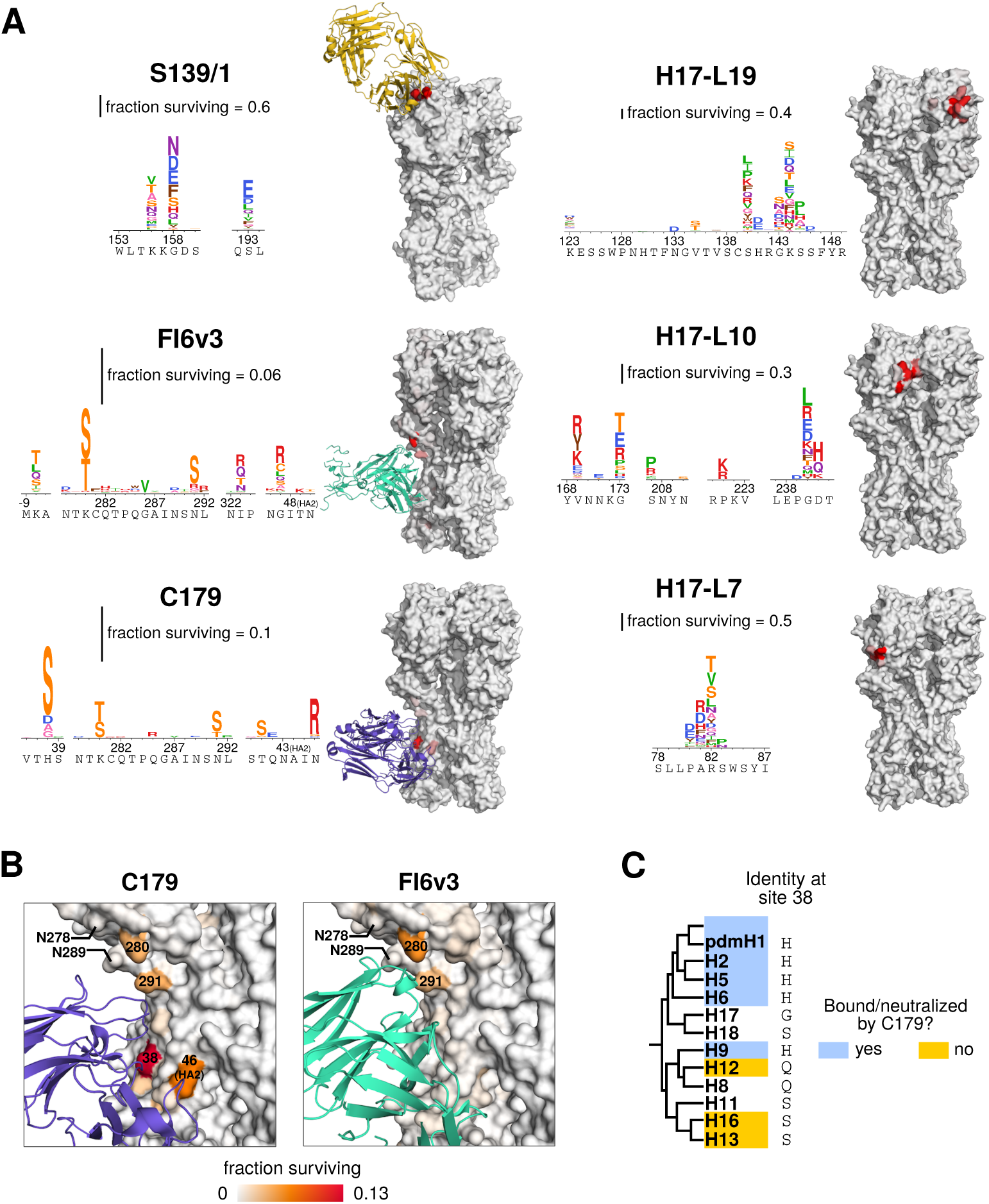
Mutations selected by broad and narrow antibodies. (A) Logo plots show sites where mutations have the largest effect. Letter heights are proportional to the excess fraction of virions with that mutation that survive antibody, as indicated by the scale bars. Structures are colored white to red by the excess fraction surviving for the largest-effect mutation at each site, with each antibody scaled separately. (B) Sites of selection from anti-stalk antibodies, with the same coloring scale for both antibodies. Selection for serine or threonine at sites 280 and 291 introduces glycosylation sites at 278 and 289, respectively. (C) Cladogram of group 1 HA subtypes. The amino acid at site 38 is indicated. Colors indicate whether a subtype has been reported in the literature to be bound or neutralized by C179.

For the S139/1 antibody that targets residues in the HA receptor-binding pocket, there are strong escape mutations at sites 156, 158, and 193 (Figure 5A; sites are in H3 numbering). These three sites fall directly in the physical binding footprint of the antibody^27^, and are the same three sites where previous work has selected escape mutants in H1, H2, and H3 HAs^41^. Our data show that numerous different amino-acid mutations at each site confer neutralization resistance. The mutation with the largest effect, G158N, introduces an N-linked glycosylation motif.

Although the anti-stalk antibodies C179 and FI6v3 only select mutations with small effects, these mutations almost all fall in or near the physical binding footprints of the antibodies (Figure 5A). The two antibodies have similar epitopes and angles of approach^50^, and they select identical mutations at several sites (Figure 5B). The three largest-effect mutations for FI6v3 (K280S, K280T, and N291S) all introduce glycosylation motifs near the epitope, and all three mutations have similar magnitude antigenic effects in both FI6v3 and C179.

However, C179 selects several mutations that do not have any apparent effect on FI6v3 (Figure 5A, Figure S6). The most notable of these C179-specific mutations are at site 38. The additional breadth of FI6v3 over antibodies such as C179 that neutralize only group 1 HAs is because FI6v3 can accommodate a glycan on the asparagine at site 38 that is present in group 2 HAs^26,24,25^. However, the H38S mutation that has the largest effect on C179 resistance in our experiments does not introduce a glycosylation motif, showing that there are also other ways to escape anti-stalk antibodies at this site. Interestingly, group 1 HA subtypes that are susceptible to C179 tend to possess a histidine at site 38, but subtypes that are not bound or neutralized by C179 often possess a serine (Figure 5C).

The FI6v3 antibody also weakly selects several mutations at residue −8, which is part of HA’s signal peptide (Figure 5A). This signal peptide is cleaved from the mature HA protein^56,57^, although mutations at this site can affect HA’s expression level^58^, which might conceivably affect HA density on virions and subsequently antibody neutralization^26,54^.

### Neutralization assays validate the mutational antigenic profiling

Do the mutations identified in our mutational antigenic profiling actually have the expected effect on antibody neutralization? We have previously validated many of the large-effect antigenic mutations selected by the narrow antibodies H17-L19, H17-L10, and H17-L7^47^. However, the mutations selected by the broad anti-stalk antibodies have much smaller effects in our mutational antigenic profiling—especially for the broadest antibody, FI6v3. We therefore tested some of these FI6v3-selected mutations using neutralization assays on individual viral mutants.

Figure 6 shows that the mutational antigenic profiling is highly predictive of the results of the neutralization assays, even for small-effect mutations. As discussed in the previous section, the three mutations most strongly selected by FI6v3 introduce glycosylation motifs at sites 278-280 or 289-291 (Figure 5A,B). We created viruses carrying each of these mutations (K280S, K280T, and N291S) and validated that all three modestly but significantly increased resistance to FI6v3 (Figure 6A, Figure S9). As a control, we also validated that a mutation at one of these sites (K280A) that does *not* have an effect in our mutational antigenic profiling does not significantly shift the neutralization curve (Figure 6A, Figure S9).

**Figure 6:**
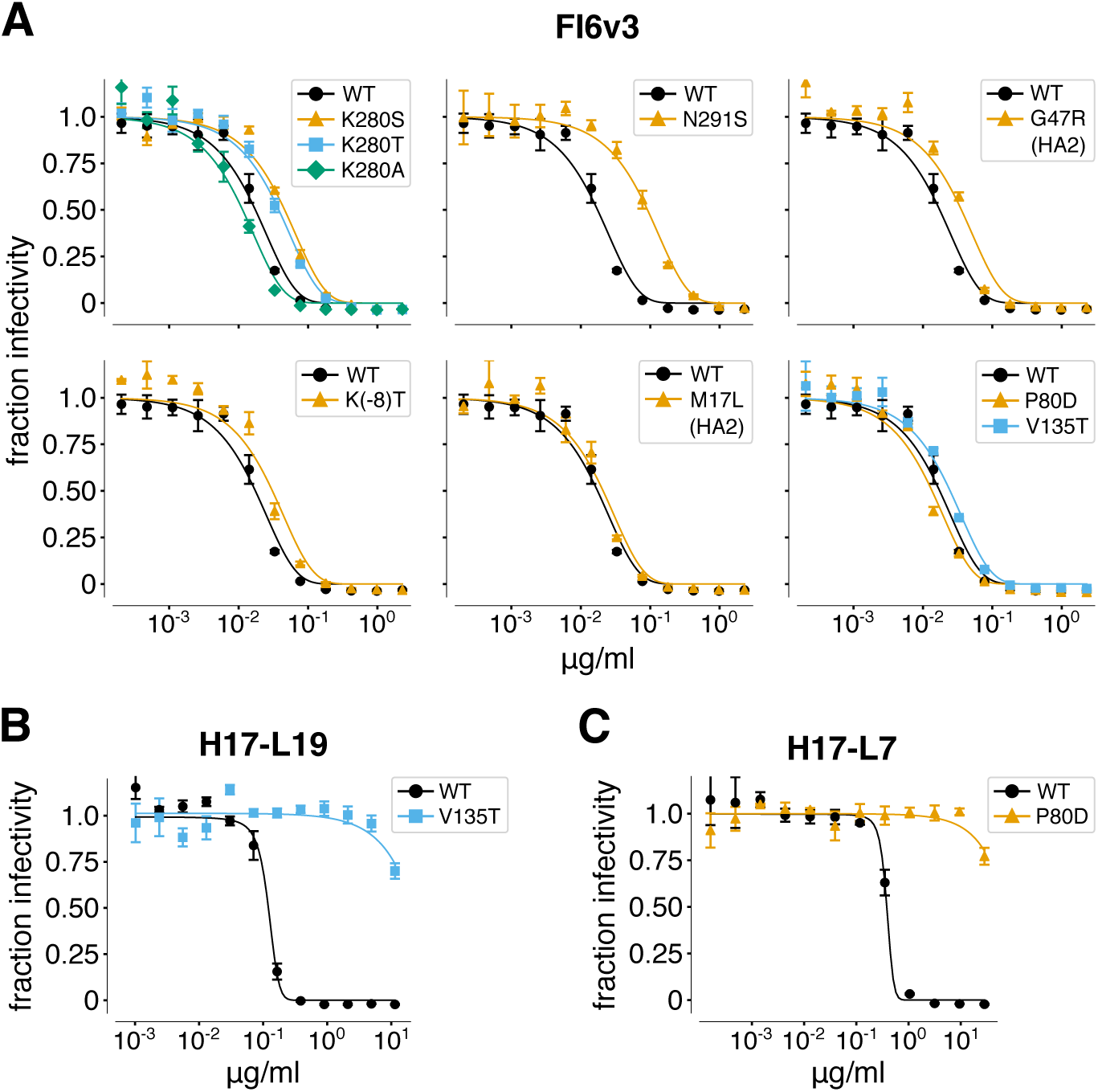
The mutations selected by FI6v3 increase neutralization resistance, but the effects are small. (A) Neutralization curves of individual viral mutants with FI6v3. The mutations K280S, K280T, N291S, G47R (HA2), and K(-8)T are all expected to increase neutralization resistance based on the mutational antigenic profiling (Figure 5A), whereas K280A, M17L (HA2), P80D, and V135T are *not* expected to affect neutralization (Figure S6). All neutralization curves in this panel were performed in triplicate on the same day. This panel shows the average of the replicates; Figure S9 shows the curves for each replicate individually and performs statistical testing of whether the IC50s for mutants are significantly different than for wildtype. (B), (C) In contrast to FI6v3, mutations selected by narrow antibodies have very large effects on neutralization. Shown are neutralization curves for representative escape mutants from H17-L19 and H17-L7 taken from Doud et al^47^.

Our mutational antigenic profiling also identified several non-glycosylation-motif mutations that were selected by FI6v3. We validated that one of these mutations, G47R in the HA2 chain, significantly increased neutralization resistance (Figure 6A, Figure S9)—although as predicted by the mutational antigenic profiling, the magnitude of the effect was small. The most unexpected mutations identified in our muta-tional antigenic profiling were at site −8 in the signal peptide. We tested one of these mutations, K(-8)T, and it did lead to a very slight increase in neutralization resistance (Figure 6A, Figure S9)—although despite the significance testing in Figure S9, we remain circumspect about the magnitude of this effect relative to the noise in our neutralization assays. As controls, we also tested three mutations (P80D and V135T, which are escape mutations for H17-L7 and H17-L19, and M17L in HA2) that did *not* have substantial effects in the mutational antigenic profiling, and confirmed that none of them significantly affected neutralization resistance (Figure 6A, Figure S9).

A notable aspect of these validation experiments is the very small effect sizes of the identified mutations on neutralization by FI6v3. Antigenic mutations selected by strain-specific antibodies to HA generally increase the concentration of antibody needed to neutralize the virus by orders of magnitude. Neutralization curves for such large-effect escape mutants are in Figure 6B,C. Although there are no such large-effect single mutations that escape FI6v3 or C179, the results in Figure 6A show that we can still use mutational antigenic profiling to identify mutations that have small but measurable effects on resistance to these antibodies.

### Limited HA mutational tolerance does not fully explain the lack of strong escape mutations from the anti-stalk antibodies

Why are there no large-effect escape mutations from the anti-stalk antibodies? One possibility is that all HA sites in the antibody-binding footprint are intolerant of mutations, meaning that viruses with mutations at these sites cannot replicate and so are not present in our mutant virus libraries. Another possibility is that mutations are tolerated at some HA sites in the antibody footprint, but that the binding energetics are distributed across sites in such a way that none of these tolerated mutations strongly affect neutralization.

We can examine these possibilities using deep mutational scanning data that measures the tolerance of HA for each possible amino-acid mutation. Specifically, we have previously selected our A/WSN/1933 virus HA mutant libraries for variants that can replicate in cell culture, and then used deep sequencing to estimate the preference of each site in HA for each possible amino acid^55^. Figure 7 shows these amino-acid preferences for all sites in HA within 4 angstroms of each broad antibody, with the antigenic effects of the mutations overlaid. Although some HA sites in the antibody footprints strongly prefer a single amino acid, for all antibodies there are also footprint sites that tolerate a fairly wide range of amino acids. In most cases the mutations selected by the antibodies occur at these mutationally tolerant sites. However, there are exceptions—for instance, the H38S mutation selected by C179 is rather disfavored with respect to viral growth, but has a large enough antigenic effect to still be detected in our mutational antigenic profiling.

**Figure 7:**
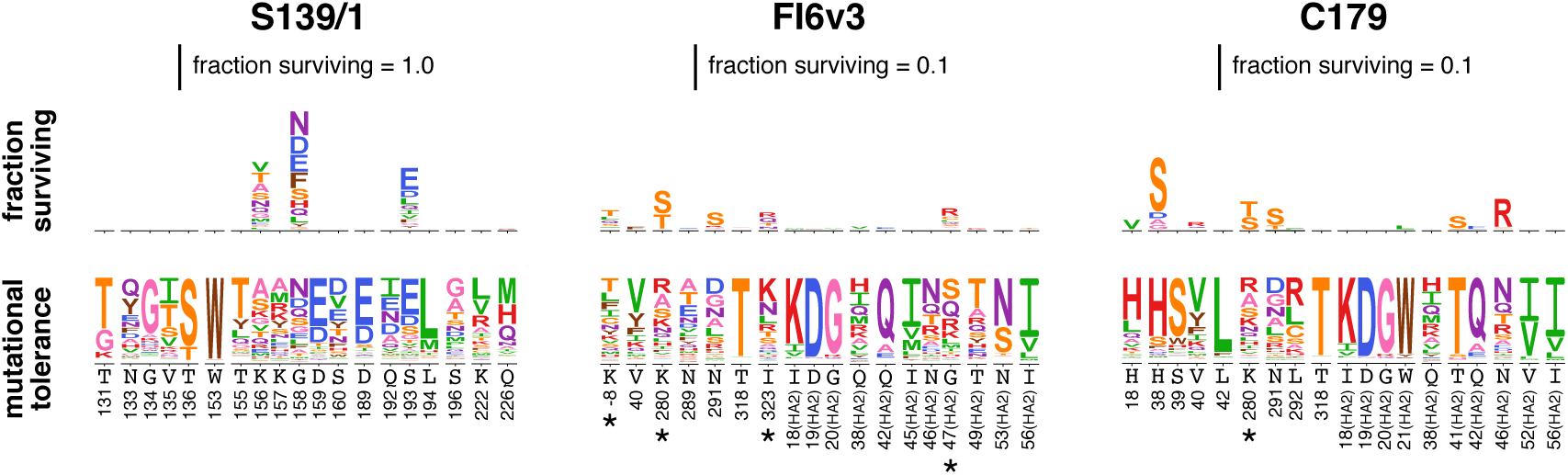
Mutational tolerance of HA sites in the antibody-binding footprints. These plots show all HA sites within 4 angstroms of the antibody in the crystal structure, plus any additional sites (marked with a *) where we identified antigenic mutations. The logo plots at bottom show the preference of each HA site for each amino acid under selection for viral replication as measured by Doud and Bloom^55^. For instance, site 153 only tolerates tryp-tophan, so W occupies the entire height of the preference logo stack. In contrast, site 156 tolerates many amino acids, all of which contribute to the height of the preference logo stack. Above the preference logo stacks are logo plots showing the excess fraction surviving antibody treatment as measured in the current study. Note that scale for these antigenic effects is 10× smaller for FI6v3 and C179 than for S139/1.

The data in Figure 7 show that the lack of large-effect escape mutants from FI6v3 and C179 is not entirely due to the mutational intolerance of HA sites in the antibody-binding footprints. Some HA sites in each antibody footprint are fairly mutationally tolerant, and contain a range of mutations in the viral libraries used in our antibody selections. However, our mutational antigenic profiling shows that only a fraction of mutations at a fraction of these sites actually affect antibody neutralization. This finding is reminiscent of prior work showing that the binding energetics at protein-protein interfaces can be asymmetrically distributed across sites^59,60,61^. The broad anti-stalk antibodies therefore appear to both mostly target mutationally intolerant sites and distribute their binding energetics in such a way that altering the mutationally tolerant HA sites has relatively little effect on neutralization.

## DISCUSSION

We have quantified how all single amino-acid mutations to an H1 influenza virus HA affect neutralization by a collection of broad and narrow antibodies. Our results show that the virus’s inherent evolutionary capacity for escape via point mutations differs across antibodies. Interestingly, antibody breadth is not always an indicator of the difficulty of viral escape. As expected, single amino-acid mutations can make the virus completely resistant to narrow strain-specific antibodies against HA’s globular head. However, such mutations can also enable the virus to escape the broad S139/1 antibody targeting residues in HA’s receptor-binding pocket, despite the fact that this antibody neutralizes multiple subtypes. But no single mutation has a comparably large effect on neutralization by two broad antibodies targeting HA’s stalk, FI6v3 and C179. Therefore, these anti-stalk antibodies are quantifiably more difficult for the virus to escape.

Although there are no large-effect escape mutations from the broad anti-stalk antibodies, there are mutations that more modestly affect neutralization. This finding emphasizes the importance of identifying antigenic mutations in a way that accounts for effect sizes. The classic approach for selecting escape mutations involves treating a virus stock with antibody at a concentration that completely neutralizes wildtype, and looking for viral mutants that survive this treatment ^16,17^. There are no such single mutations for the H1 HA and broad anti-stalk antibodies tested here, since no mutations shift the neutralization curve enough to enable survival at antibody concentrations that fully neutralize wildtype. However, our approach shows that there are mutations that have more modest (*<*10-fold) effects on neutralization by even the broadest antibody Interestingly most previous studies^42,43,44,45^ that have reported selecting single mutations with large effects (»10-fold) on neutralization by anti-stalk antibodies have used group 2 (e.g., H3 or H7) HAs rather than group 1 HAs like the one used in our work. When interpreting the magnitude of these effects, it is important to note that our experiments only measure how mutations affect neutralization, and not how they affect Fc-mediated functions that are responsible for much of the *in vivo* protection afforded by anti-stalk antibodies^62,63^.

Another important caveat is that our experiments examine *single* amino-acid mutations to the HA from one influenza virus strain. The protein evolution literature is full of examples of epistatic interactions that enable multiple mutations to access phenotypes not accessible by single mutations^64,65,66^. Such epistasis is relevant to HAs evolution. For instance, work by Das et al^67^ suggests that the sequential accumulation of mutations can shift the spectrum of available antibody-escape mutations. Wu et al^68^ have used deep mutational scanning to directly demonstrate that rampant epistasis enables HAs receptor-binding pocket to accommodate combinations of individually deleterious mutations, some of which affect sensitivity to antibodies. Therefore, our work does not imply any absolute limits on the possibilities for antibody escape when evolution is given sufficient time to explore combinations of mutations. However, single mutations are the most accessible form of genetic variation, and much of influenza virus’s natural antigenic drift involves individual mutations that reduce sensitivity to immunodominant antibody specificities^16,17,18,19,20,21^. Quantifying the antigenic effects of all such mutations therefore provides a relevant measure of ease of viral antibody escape.

A major rationale for studying broadly neutralizing antibodies is that they are hoped to be more resistant to viral evolutionary escape than the antibodies that dominate natural immune responses to influenza virus^32,33^. We have used a new approach to quantify the extent to which this is actually true, and shown that neutralization of an H1 virus by broad anti-stalk antibodies is indeed more-although certainly not completely—resistant to erosion by viral point mutations. Going forward, we suggest that completely mapping viral escape mutations will be a useful complement to more traditional techniques that simply characterize the breadth of anti-viral antibodies against circulating strains.

## METHODS

### Antibodies

C179 IgG was purchased from Takara Bio Inc (Catalog #M145). FI6v3 was purified from 293F cells transduced with a lentiviral vector encoding a commercially synthesized gene for the IgG form of the antibody, with the heavy and light chains reverse-translated from the protein sequence in the PDB structure 3ZTN^26^ as described previously^69^. Genes encoding S139/1 in IgG form were were reverse-translated from the protein sequence in PDB structure 4GMS^27^, and used to express and purify protein by the Fred Hutchinson Cancer Research Center protein expression core.

### Neutralization assays

We performed neutralization assays using influenza viruses that carried GFP in the PB1 segment. These PB1flank-eGFP were generated in co-cultures of 293T-CMV-PB1 and MDCK-SIAT1-CMV-PB1 cells as described previously^70^, using the standard bi-directional pHW181-PB2,…, pHW188-NS reverse-genetics plasmids^71^ for all genes *except* PB1, plus the pHH-PB1flank-eGFP plasmid^70^. Each mutant was generated by repeating this process using a version of the pHW184-HA plasmid that had been engineered by site-directed mutagenesis to carry the indicated mutation. The neutralization assays themselves were performed by using a plate reader to quantify the GFP signal produced by MDCK-SIAT1-CMV-PB1 cells infected by PB1flank-eGFP virus that had been incubated with the indicated antibody concentration as described previously^72^. All neutralization curves in Figure 6A represent the mean and standard deviation of three measurements, with the individual replicates shown in Figure S9. All the neutralization assays for FI6v3 were performed on the same day to eliminate batch effects, with each replicate involving independent serial dilution of the antibody in a separate column of a 96-well plate.

### H3 sequence numbering

Unless otherwise indicated, all residues are numbered in the H3 numbering scheme, with the signal peptide in negative numbers, the HA1 subunit as plain numbers, and the HA2 subunit denoted with “(HA2)”. The conversion between sequential numbering of the A/WSN/1933 HA and the H3 numbering scheme was performed using the Python script available at https://github.com/jbloomlab/HA_numbering. File S1 gives the numbering conversion.

### Inference of HA phylogenetic tree

To infer the phylogenetic tree in Figure 2, we downloaded one HA sequence per subtype from the Influenza Research Database^73^, inferred the phylogenetic tree using RaxML^74^ with a GTR model, and visualized the tree using FigTree (http://tree.bio.ed.ac.uk/software/figtree/). The HA sequences used are in File S2. In Figure 2, we indicate which HAs each antibody has been reported to bind or neutralize^41,27,51,50^,. Among broad antibodies, S139/1 has not been tested against H8 and H11; C179 has not been tested against H8 and H11; and no antibodies have been tested against H17 and H18. The narrow H17-L19, H17-L10, and H17-L7 antibodies have not been tested against any other subtypes—however, since these antibodies have a very limited range even among H1 HAs^52^, we assume that they do not bind other subtypes.

For the cladogram in Figure 5C, the amino-acid identities at site 38 are from the strains tested against C179 by by Dreyfus et al^50^. For subtypes not tested, the amino-acid identity reported is that in the strain for that subtype in File S2.

### Mutant virus libraries

The mutant virus libraries are those described in Doud and Bloom^55^, and were produced in full biological triplicate. Briefly, these libraries were generated by using codon mutagenesis^75^ to introduce random codon mutations into plasmid-encoded HA, and then using a helper-virus strategy that avoids the bottlenecks associated with standard influenza reverse genetics to create the virus libraries. Although a helper virus is used to generate the libraries from plasmids, the viruses in the resulting library carry the full complement of genes and are fully infectious and replication-competent^55^. This fact is important, since the accessibility of HA epitopes can depend on virion HA density, which is often lower in pseudovirus than in fully infectious virus^26,54^. Full details of the library generation and sequencing statistics that quantify how completely each of the triplicate libraries covers the possible amino-acid mutations have been described previously^55^.

### Mutational antigenic profiling

The mutational antigenic profiling was performed as described previously^47^. Briefly, we diluted each of the virus libraries to a concentration of 10^6^ TCID_50_ per ml and incubated the virus dilutions with an equal volume of antibody at the intended concentration at 37°C for 1.5 hours. The final antibody concentrations in these mixtures are shown in Figure 3. We performed three fully independent replicates of each selection using the three replicate mutant virus libraries. In addition, we performed technical replicates (independent neutralization experiments on the *same* virus library) in some cases as indicated in Figure S1. The virus-antibody mixtures were used to infect cells, and viral RNA was extracted, reverse-transcribed, and PCR amplified as described previously^47^. In order to obtain high accuracy in the Illumina deep sequencing, we used the barcoded-subamplicon sequencing strategy described by Doud and Bloom^55^, which is a slight modification of the strategy of Wu et al^39^.

We also estimated the overall fraction of virions surviving each antibody selection. These fractions are denoted by *γ* in this paper. The average of these fractions across libraries are reported in Figure 3, and the values for each individual replicate are in Table S1. The frac-tions were estimated using qRT-PCR against the viral NP and canine GAPDH as described previously^47^. Briefly, we made duplicate 10-fold serial dilutions of each of the virus libraries to use as a standard curve of infectivity. We also performed qPCR on the cells infected with the virus-antibody mix. To estimate the fractions, we used linear regression to fit a line relating logarithm of the viral infectious dose in the standard curve to the difference in Ct values between NP and GAPDH, and then interpolated the fraction surviving for each selection from this regression.

### Analysis of deep sequencing data

The deep sequencing data were analyzed using version 2.2.1 of the dms_tools2 software package^76^, which is available at http://jbloomlab.github.io/dms_tools2. File S3 contains a Jupyter notebook that performs all steps of the analysis beginning with downloading the FASTQ files from the Sequence Read Archive. Detailed statistics about the sequencing depth and error rates are shown in this Jupyter notebook and its HTML rendering in File S4.

### Calculating the fraction of virions with each mutation that escapes antibody neutralization

In prior mutational antigenic profiling work^47,48^, we calculated the differential selection on each mutation as the logarithm of its enrichment relative to wildtype in an antibody-selected sample versus a mock-selected control. These *mutation differential selection* values are useful for the analysis of individual experiments. However, there is no natural way to compare these values across experiments with different antibodies at different concentrations, since the strength of differential selection depends on details of how the pressure is imposed. We therefore developed the new approach in this paper to quantify the antigenic effect of a mutation in units that can be compared across antibodies and concentrations.

The general principle of the calculations is illustrated in Figure 1 and discussed in the first section of the RESULTS. Here we provide details on how these calculations are performed. The deep sequencing measures the number of times that codon *x* is observed at site *r* in both the antibody-selected and mock-selected conditions. Denote these counts as 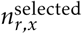 and 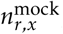, respectively. We also perform deep sequencing of a control (in this case, plasmid DNA encoding the wildtype HA gene) to estimate the sequencing error rate. Denote the counts of codon *x* at site *r* in this control as 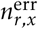. Also denote the total reads at each site *r* in each sample as 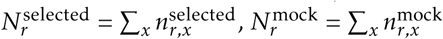, and 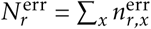.

We first estimate the rate of sequencing errors at site *r* as

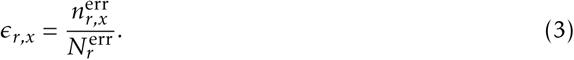

For the wildtype identity at site *r*, which we denote as wt(*r*), the value of ∈_*r,wt*(*r*)_ is the fraction of times we correctly observe the wildtype identity wt(*r*) at site *r* versus observing some spurious mutation. For all mutant identities *x* ≠ wt(*r*) at site *r*, *∈*_*r,x*_ is the fraction of times we observe the mutation *x* at site *r* when the identity is really wildtype. We ignore second-order terms where we incorrectly read one mutation as another, as such errors will be very rare as mutations themselves are rare (most codons are wildtype in most sequences).

We next adjust all of the deep sequencing codon counts in the antibody-selected and mock-selected conditions by the error control. Specifically the error-adjusted counts for the antibody-selected sample are

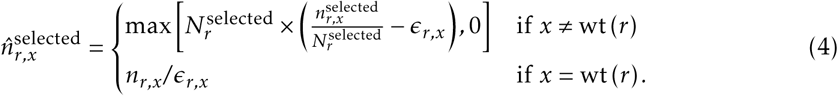

An equivalent equation is used to calculate 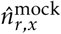. We then sum the error-adjusted codon counts for each amino acid *a*:

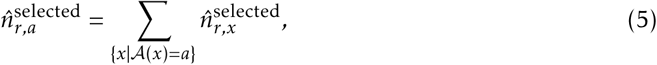

so that 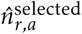 are the error-adjusted counts for the antibody-selected condition summed across all codons *x* where the encoded amino acid *A (x*) is *a*. An equivalent equation is used to calculate 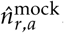.

Finally we use these error-adjusted amino-acid counts to estimate the mutation frequencies 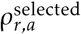 and 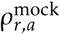 that are used in Equation 1 to calculate the fraction *F*_*r,a*_ of virions with amino acid *a* at site *r* that survive the selection. When estimating these mutation frequencies, we add a pseudocount of *P* = 5 to the lower-depth sample, and a depth-adjusted pseudocount to the higher depth sample. The rationale for adding a pseudocount is to regularize the estimates in the case of low counts. Specifically, we estimate the mutation frequencies as

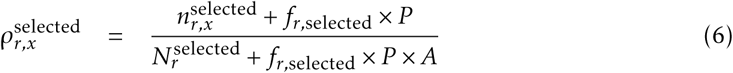

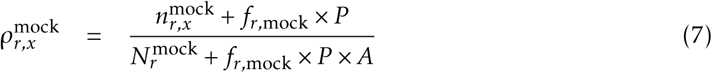

where *A* is the number of characters (e.g., 20 for amino acids), *f*_*r,selected*_ and *f*_*r,mock*_ are the pseudocount adjustment factors defined as:

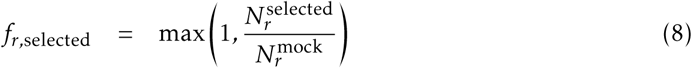

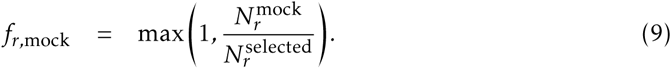

The pseudocount adjustment factors ensure that *P* is added to the counts for the lower depth sample, and a proportionally scaled-up pseudocount is added to the higher depth sample. The depth scaling is necessary to avoid systematically biasing towards higher mutation frequencies in the lower depth sample. It is these estimated mutation frequencies that are used in conjunction with *γ* (the qPCR estimated overall of virions that survive selection) to compute the fraction surviving (*F*_*r,a*_) and excess fraction surviving above the library average 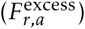 via Equations 1 and 2.

In some cases, we need to summarize the excess fraction of mutations surviving into a single number for each site, such as for plotting as a function of the site number or displaying on the crystal structure. There are 19 different 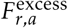 values for non-wildtype amino acids for each site. One summary statistic is the fraction surviving above the library average *averaged* over all 19 amino-acid mutations at site *r*:

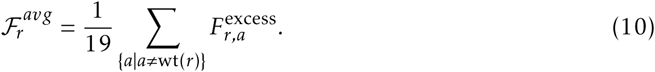

Another summary statistic is the *maximum* fraction surviving above average among all 19 amino-acid mutations at site *r*:

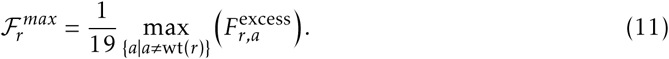

In this paper, Figures 4 and S2 show the median of excess fraction surviving taken across all biological and technical replicates at a given antibody concentration (Equation 2). The subsequent logo plots show the medians of these values taken across all concentrations for each antibody. The numerical values plotted in these logo plots are in File S5. The fraction surviving values *not* adjusted to be in excess of the library average (Equation 1) are in File S6.

Code that performs these fraction surviving analyses has been added to version 2.1.0 of the dms_tools2 software package^76^ which is available at http://jbloomlab.github.io/dms_tools2.

## Data availability and source code

Deep sequencing data are available from the Sequence Read Archive under BioSample accession SAMN05789126. Computer code that analyzes these data to generate all the results described in this paper is in File S3, and an HTML version of the analysis notebook is in File S4.

## ACKNOWLEDGMENTS

We thank Adam Dingens, Sarah Hilton, Katherine Xue, Lauren Gentles, and Jeremy Roop for helpful comments on the project and manuscript. We thank the Fred Hutchinson Cancer Research Center genomics core for performing the Illumina deep sequencing, and the protein expression core for expressing and purifying the S139/1 antibody. This work was supported by grant R01AI127893 from the NIAID of the NIH. MBD was supported in part by training grant T32AI083203 from the NIAID of the NIH. JML was supported in part by the Center for Inference and Dynamics of Infectious Diseases (CIDID), which is funded by grant U54GM111274 from the NIGMS of the NIH. The research of JDB is supported in part by a Faculty Scholar Grant from the Howard Hughes Medical Institute and the Simons Foundation. The funders had no role in study design, data collection and analysis, decision to publish, or preparation of the manuscript.

## AUTHOR CONTRIBUTIONS

MBD and JML performed the experiments. All three authors designed the project, contributed to the computer code, analyzed the data, and wrote the paper.

## Supplementary Material

**Figure S1:**
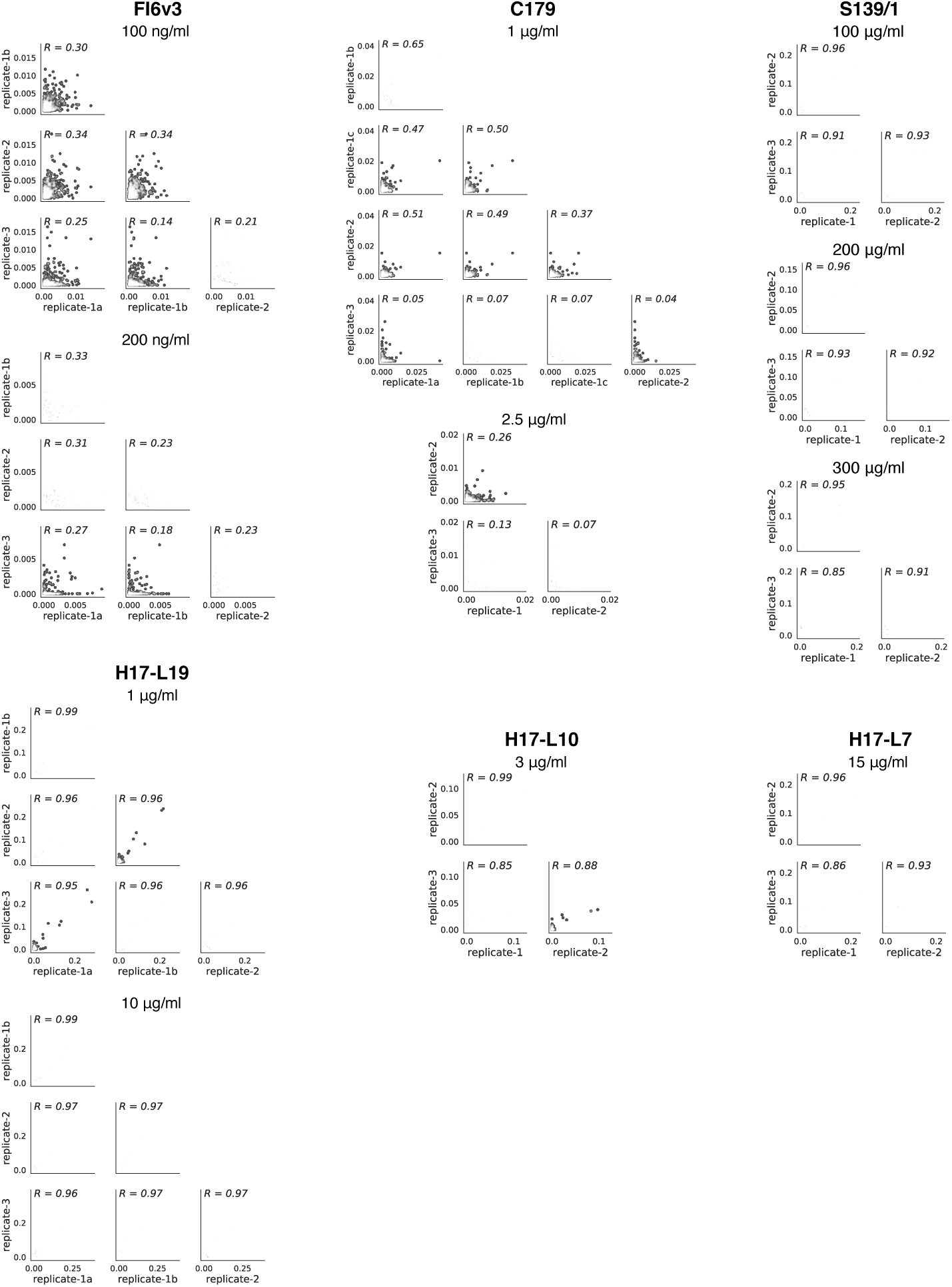
Correlations across experimental replicates. Each point represents one site in HA, and gives the fraction surviving above average across all amino-acid mutations at that site, as calculated using Equation 10. The replicates are highly correlated for antibodies with strong escape mutations (S139/1, H17-L19, H17-L10, and H17-L7), and reasonably correlated for antibodies with only weak escape mutations (FI6v3 and C179).

**Figure S2:**
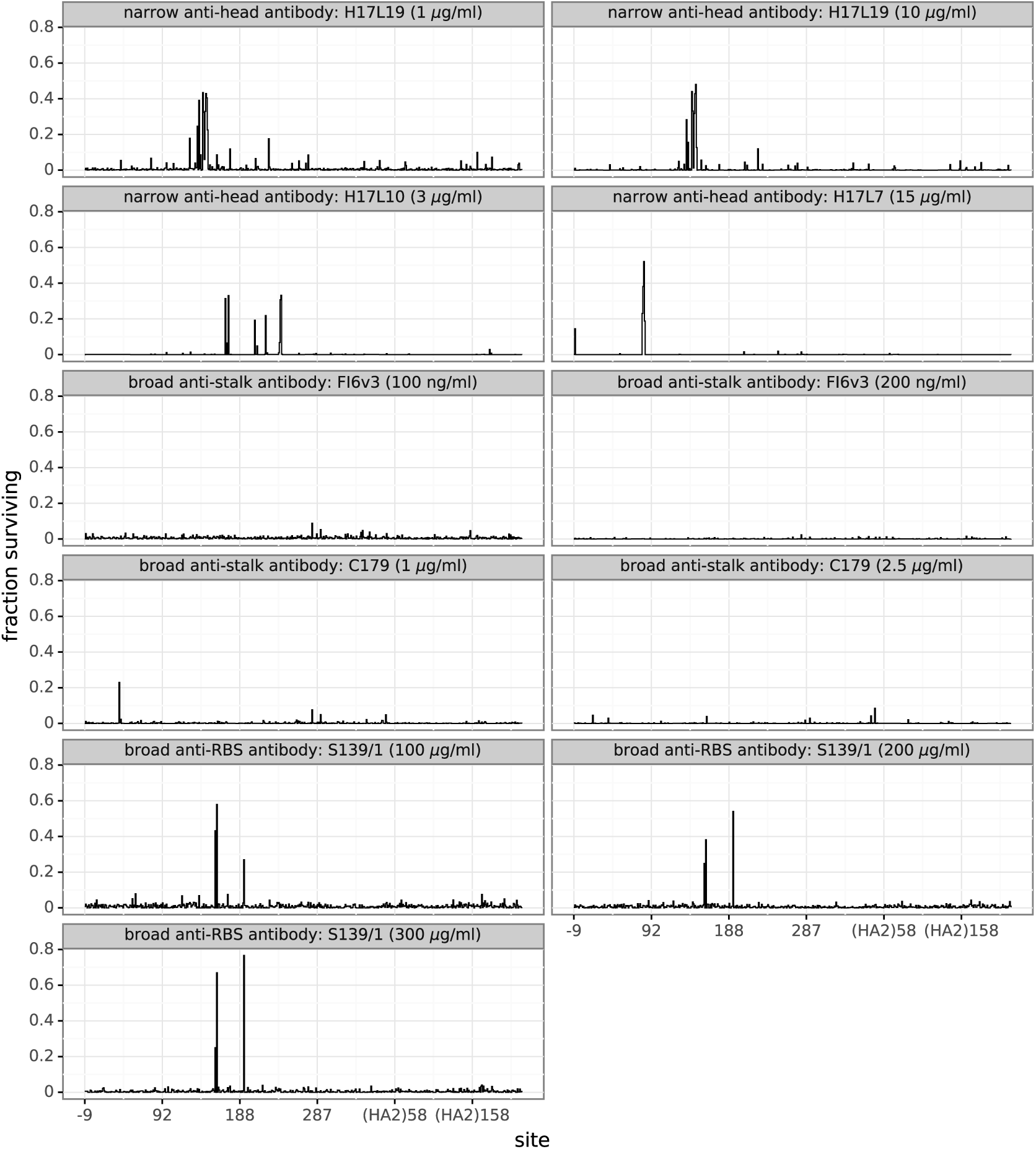
The excess fraction surviving for the single strongest escape mutation at each site. This plot differs from Figure 4 in that the height of the line indicates the excess fraction of virions that survive the antibody selection for the single strongest escape mutation at that site, rather than the average across all amino-acid mutations at that site.

**Figure S3:**
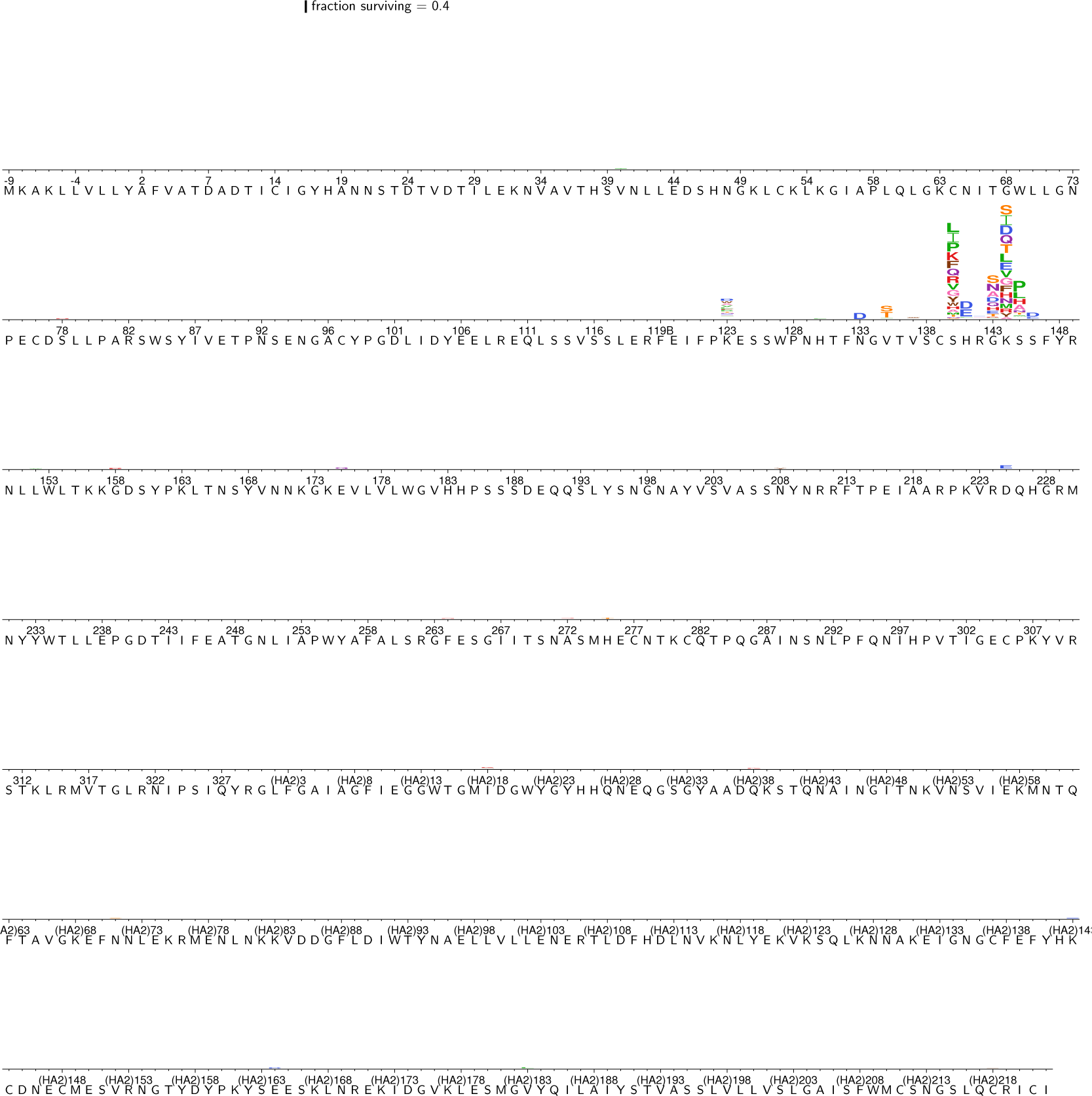
The excess fraction surviving selection with antibody H17L19 for all amino-acid mutations. The excess fraction surviving for each replicate was computed using Equation 2, then we took the median across all technical and biological replicates for each antibody concentration, and then took the medians of those values across concentrations. The height of each letter is proportional to the excess fraction surviving of virions with that mutation. The scale bar at the top of the plot relates the letter heights to the actual fractions. The sites are labeled using H3 numbering.

**Figure S4:**
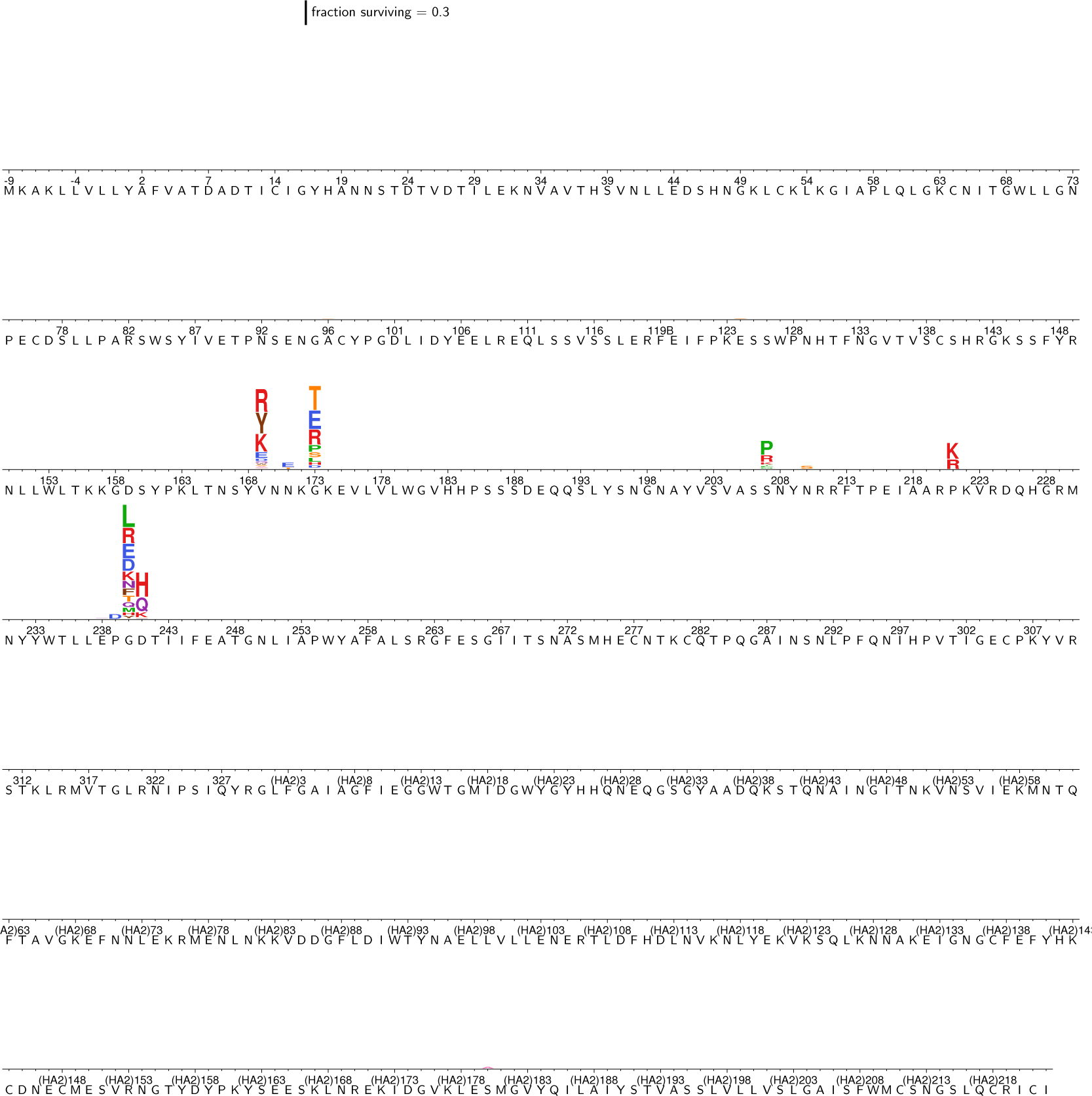
The excess fraction surviving selection with antibody H17L10 for all amino-acid mutations. The excess fraction surviving for each replicate was computed using Equation 2, then we took the median across all technical and biological replicates for each antibody concentration, and then took the medians of those values across concentrations. The height of each letter is proportional to the excess fraction surviving of virions with that mutation. The scale bar at the top of the plot relates the letter heights to the actual fractions. The sites are labeled using H3 numbering.

**Figure S5:**
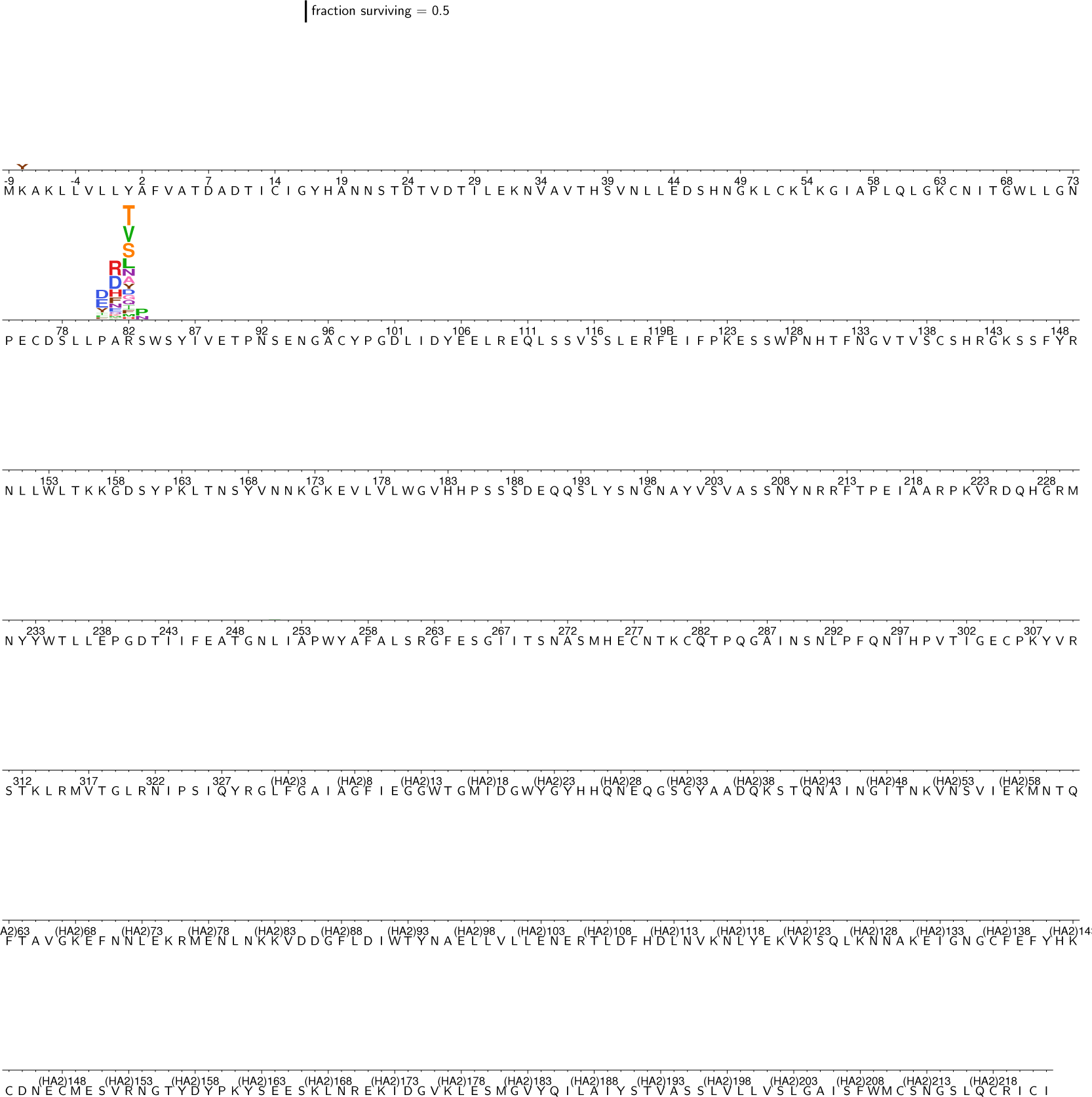
The excess fraction surviving selection with antibody H17L7 for all amino-acid mutations. The excess fraction surviving for each replicate was computed using Equation 2, then we took the median across all technical and biological replicates for each antibody concentration, and then took the medians of those values across concentrations. The height of each letter is proportional to the excess fraction surviving of virions with that mutation. The scale bar at the top of the plot relates the letter heights to the actual fractions. The sites are labeled using H3 numbering.

**Figure S6:**
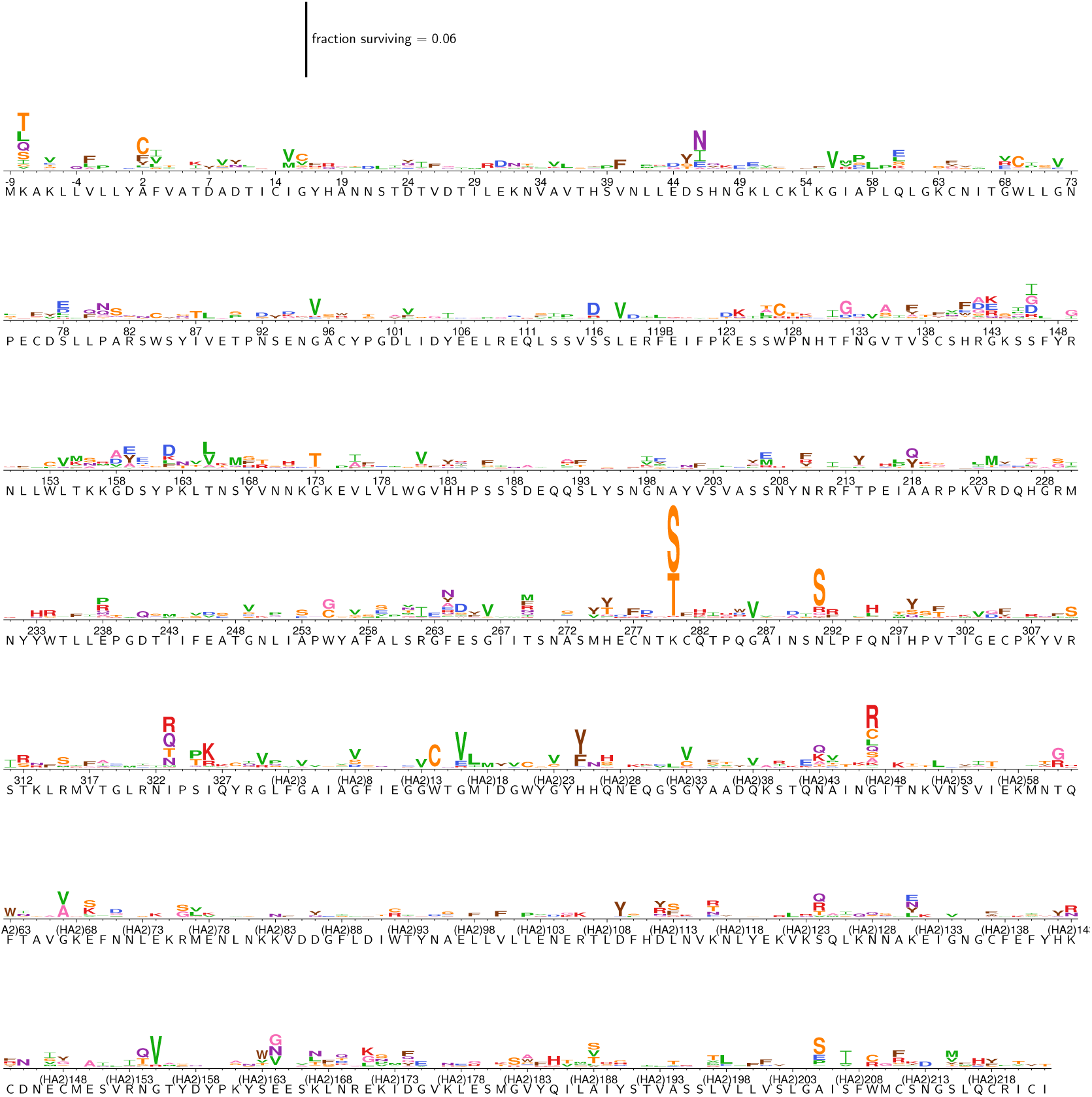
The excess fraction surviving selection with antibody FI6v3 for all amino-acid mutations. The excess fraction surviving for each replicate was computed using Equation 2, then we took the median across all technical and biological replicates for each antibody concentration, and then took the medians of those values across concentrations. The height of each letter is proportional to the excess fraction surviving of virions with that mutation. The scale bar at the top of the plot relates the letter heights to the actual fractions. The sites are labeled using H3 numbering.

**Figure S7:**
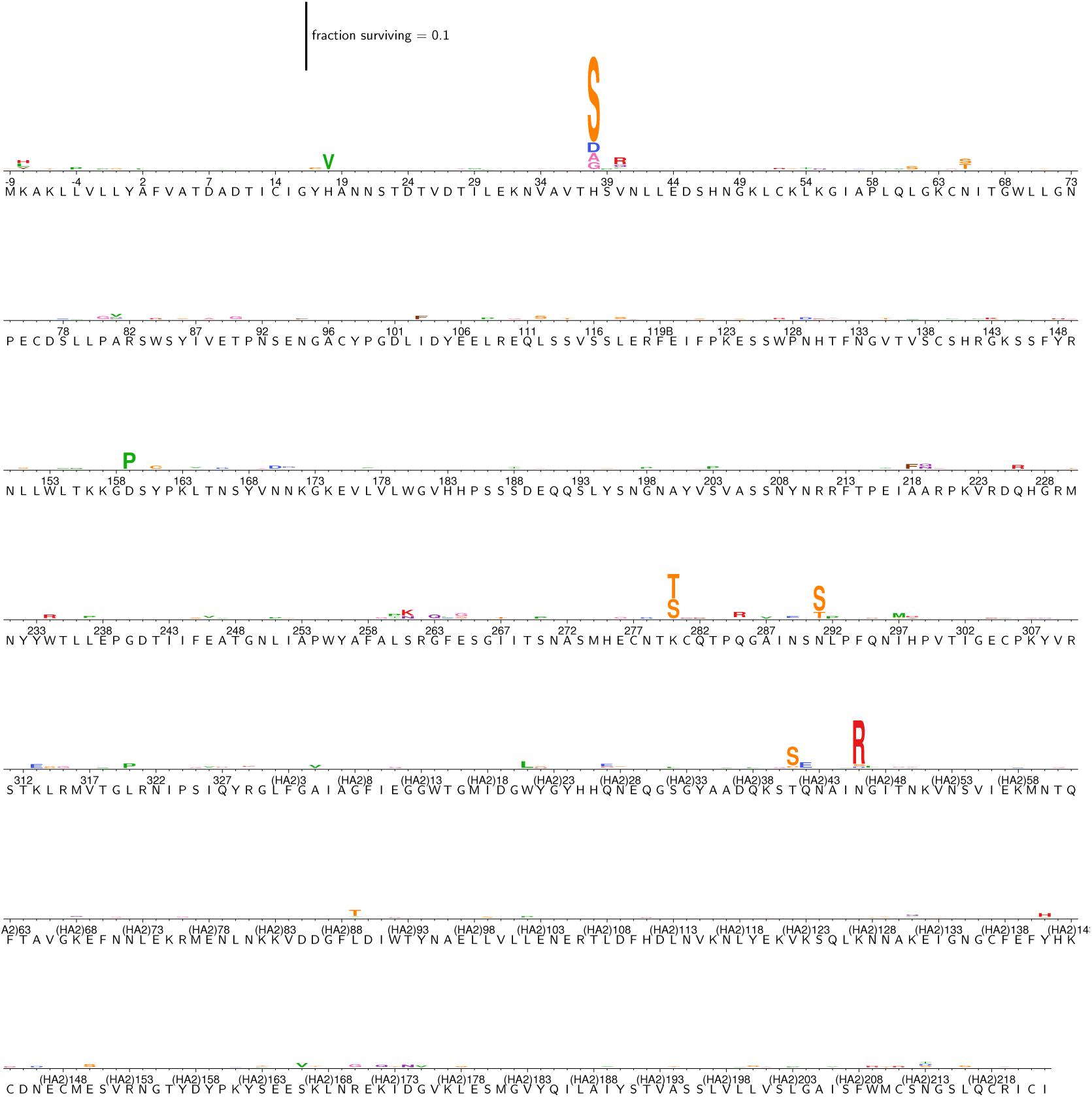
The excess fraction surviving selection with antibody C179 for all amino-acid mutations. The excess fraction surviving for each replicate was computed using Equation 2, then we took the median across all technical and biological replicates for each antibody concentration, and then took the medians of those values across concentrations. The height of each letter is proportional to the excess fraction surviving of virions with that mutation. The scale bar at the top of the plot relates the letter heights to the actual fractions. The sites are labeled using H3 numbering.

**Figure S8:**
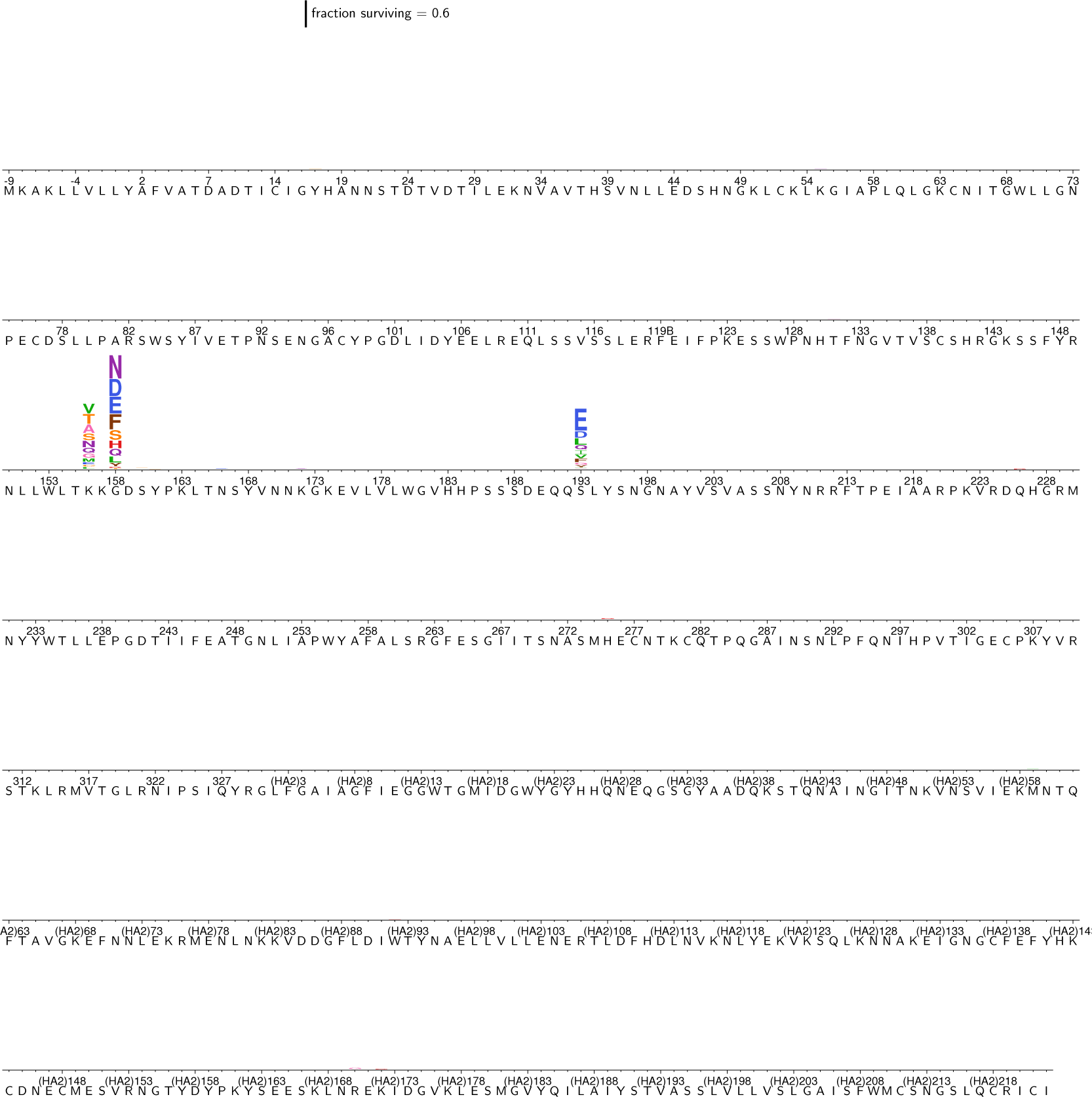
The excess fraction surviving selection with antibody S139/1 for all amino-acid mutations. The excess fraction surviving for each replicate was computed using Equation 2, then we took the median across all technical and biological replicates for each antibody concentration, and then took the medians of those values across concentrations. The height of each letter is proportional to the excess fraction surviving of virions with that mutation. The scale bar at the top of the plot relates the letter heights to the actual fractions. The sites are labeled using H3 numbering.

**Figure S9:**
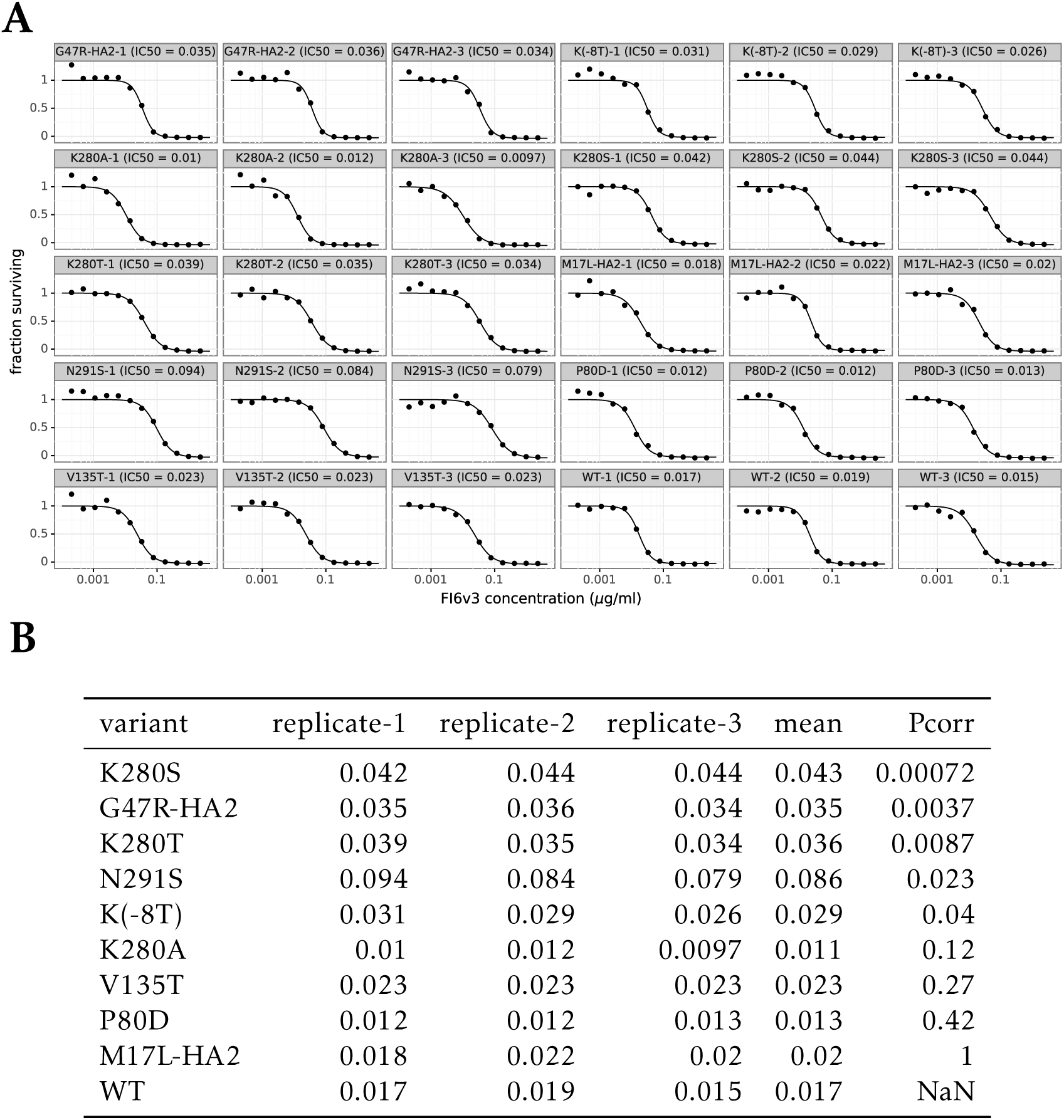
Replicates of the FI6v3 neutralization curves in Figure 6A. The neutralization assays were performed in triplicate for all nine mutants and wildtype. Figure 6A shows the *average* of those replicates. (A) The neutralization data for each replicate shown individually, with IC50 values fit using a four-parameter logistic curve with the top value constrained to one (see https://jbloomlab.github.io/dms_tools2/dms_tools2.neutcurve.html for the code used for the fitting.) (B) Table of the IC50 values for each replicate. We used an unpaired Student’s t-test with unequal variances to test the null hypothesis that each mutant had an IC50 indistinguishable from wildtype. We then used Bonferroni’s method to correct the *P*-values for multiple testing, and report these corrected values.

**Table S1:**
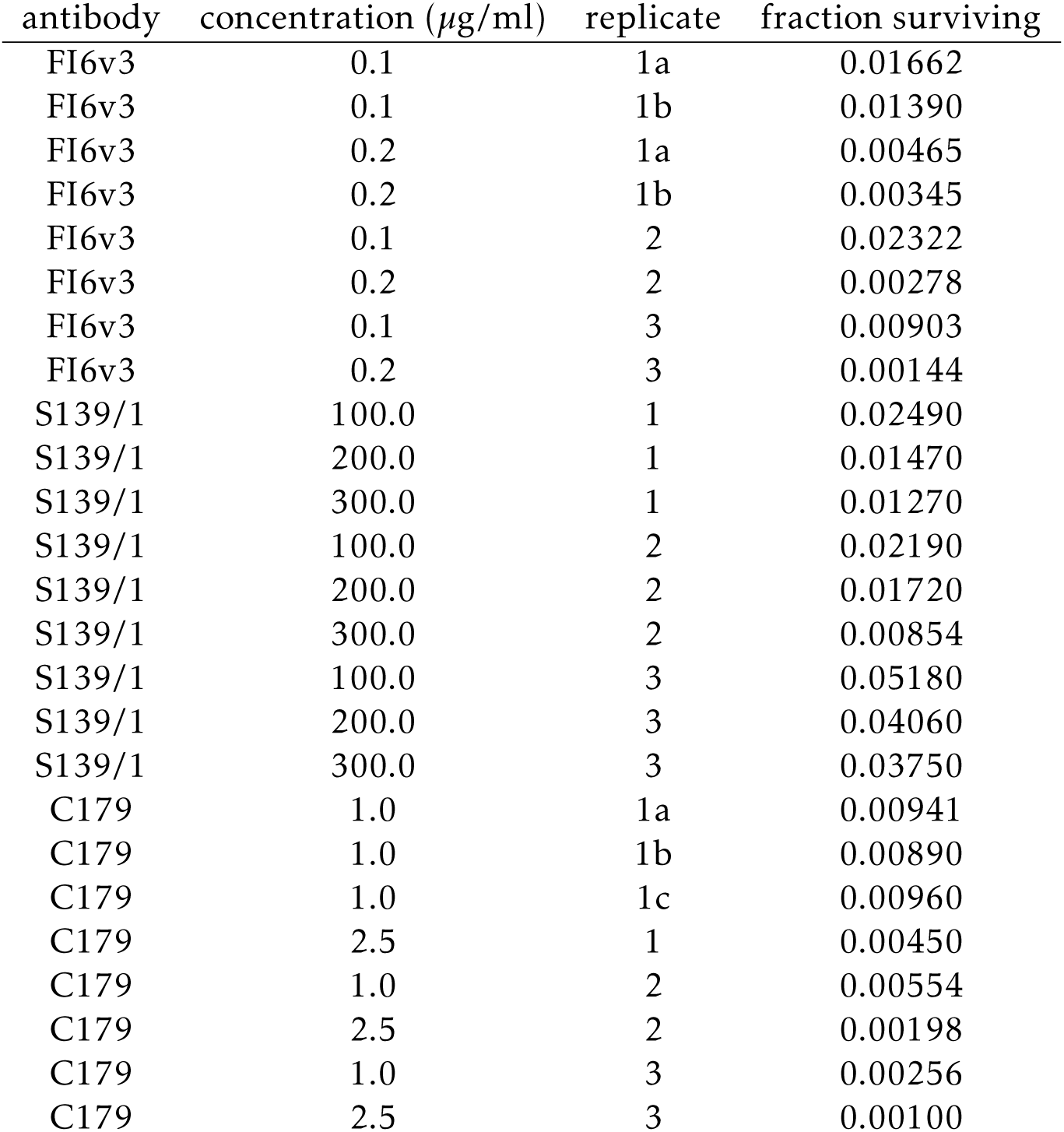
The total fraction of virions surviving each antibody treatment at each concentration as estimated by qPCR. These are the quantities referred to as *γ*. This table shows the values for the broad antibodies; values for the narrow H17-L17, H17-L10, and H17-L7 antibodies have been reported previously^47^.

**File S1: Conversion from sequential numbering of the A/WSN/1933 HA to H3 numbering**. In this CSV file, the *original* column gives the residue number in sequential (1, 2,…) numbering of the A/WSN/1933 HA, and the *new* column gives the residue number in H3 numbering.

**File S2: Sequences used to infer the tree for all HA subtypes**. This FASTA file gives the HA sequences used to infer the tree of subtypes in Figure 2.

**File S3: Computer code and data for the analysis of the mutational antigenic profiling data**. The code in this ZIP file performs the entire computational analysis beginning with downloading the FASTQ files from the Sequence Read Archive. The ZIP file contains a README file that explains the contents in detail. The actual analysis is performed by the Jupyter notebook analysis_notebook.ipynb, which includes embedded plots summarizing key statistics and results. An HTML version of this notebook is also included as File S3.

**File S4: HTML version of the analysis notebook**. This file is an HTML rendering of the Jupyter notebook in File S3. It contains detailed plots for all aspects of the deep sequencing data and its analysis.

**File S5: The excess fraction surviving for each mutation for each antibody**. This file is a ZIP of CSV files giving the numerical values plotted in the logo plots. These are median excess fraction surviving taken first across replicates and then across antibody concentrations. See Equation 2.

**File S6: The fraction surviving for each mutation for each antibody**. This file differs from File S5 only in that the values are *not* adjusted to be in excess of the library average (e.g., they are from Equation 1 rather than Equation 2).

